# Coexistence with *Pseudomonas aeruginosa* alters *Staphylococcus aureus* transcriptome, antibiotic resistance and internalization into epithelial cells

**DOI:** 10.1101/775684

**Authors:** Paul Briaud, Laura Camus, Sylvère Bastien, Anne Doléans-Jordheim, François Vandenesch, Karen Moreau

## Abstract

Cystic fibrosis (CF) is the most common life-threatening genetic disease among Caucasians. CF patients suffer from chronic lung infections due to the presence of thick mucus, caused by *cftr* gene dysfunction. The two most commonly found bacteria in the mucus of CF patients are *Staphylococcus aureus* and *Pseudomonas aeruginosa*. It is well known that early-infecting *P. aeruginosa* strains produce anti-staphylococcal compounds and inhibit *S. aureus* growth. More recently, it has been shown that late-infecting *P. aeruginosa* strains develop commensal-like/coexistence interaction with *S. aureus*. The aim of this study was to decipher the impact of *P. aeruginosa* strains on *S. aureus*. RNA sequencing analysis showed 77 genes were specifically dysregulated in the context of competition and 140 genes in the context of coexistence in the presence of *P. aeruginosa*. In coexistence, genes encoding virulence factors and proteins involved in carbohydrates, lipids, nucleotides and amino acids metabolism were downregulated. On the contrary, several transporter family encoding genes were upregulated. In particular, several antibiotic pumps belonging to the Nor family were upregulated: *tet38*, *norA* and *norC*, leading to an increase in antibiotic resistance of *S. aureus* when exposed to tetracycline and ciprofloxacin and an enhanced internalization rate within epithelial pulmonary cells. This study shows that coexistence with *P. aeruginosa* affects the *S. aureus* transcriptome and virulence.

## INTRODUCTION

Most microorganisms are frequently embedded within communities of mixed species where different microbial interactions can occur between individual species. In the case of infection, these interactions between species can influence pathogenic behavior such as virulence, biofilm formation and antibiotic tolerance ^1–4^.

One of the most well-known examples of pathologies in which many bacterial interactions are described are lung diseases occurring during Cystic Fibrosis (CF). The airways of CF patients are colonized by multiple microorganisms whose prevalence varies with the age of the patients. Among them, *Staphylococcus aureus* and *Pseudomonas aeruginosa* are the most prevalent pathogens and are acquired in subsequent order. The typical pattern of chronic infection establishment begins with the early acquisition of *S. aureus*, (60% prevalence among children aged <2 years and the highest prevalence in children of 11-17 years (80%)), while prevalence slowly declines in adults (50%) ^5^. In contrast, infections by *P. aeruginosa* occur later with the highest prevalence in adults (70% among 35-44-year-old patients). Although these bacteria seem to succeed one another, they are not mutually exclusive since patients are frequently diagnosed as being co-infected by *S. aureus* and *P. aeruginosa* (from 35% to 50%) ^6,7^.

While *P. aeruginosa* is recognized as the leading cause of lung function decline, the significance of *S. aureus* in the course of CF disease is still being debated. It has been shown that one of the risk factors for initial *P. aeruginosa* airway infection includes *S. aureus* pre-colonization ^8–10^. However, the impact of coinfection by the two pathogens on the evolution of the disease remains unclear ^11–13^.

*S. aureus* and *P. aeruginosa* have been identified in the same lobe of CF lungs ^14,15^, suggesting that both pathogens are present in the same niche and can in fact interact *in vivo*. Interactions have been widely studied and it is commonly admitted that *P. aeruginosa* outcompetes *S. aureus*. Different mechanisms have been described^16^: for example, *P. aeruginosa* secreted products can inhibit the growth or lyse *S. aureus* as well as induce epithelial cells to kill *S. aureus* and other Gram-positive bacteria ^8,17,18^.

However, these interactions can evolve during chronic colonization. Indeed, *P. aeruginosa* strains isolated from early infection outcompete *S. aureus*, as previously described, while strains isolated from chronic infection are less aggressive and can be co-cultivated with *S. aureus* ^19,20^. Furthermore, *P. aeruginosa* isolates from mono-infected patients are more competitive towards *S. aureus* than isolates from coinfected patients ^21^.

In contrast to antagonistic interactions, nothing is known about the effects of *P. aeruginosa* and *S. aureus* interactions in this context of coexisting bacteria within the same infectious niche. Using a transcriptomic approach, we analyzed how co-cultivation with non-competitive *P. aeruginosa* altered *S. aureus* gene expression, especially genes encoding Nor family efflux pumps. In the presence of *P. aeruginosa*, over-expression of these genes increased *S. aureus* antibiotic tolerance and the rate of internalization into epithelial cells, two key determinants of chronic infection.

## RESULTS

### Coexistence interaction involves more than half of the *S. aureus* and *P. aeruginosa* isolates from co-infected CF patients

Two types of interactions between *S. aureus* and *P. aeruginosa* could be observed with CF patient isolates: the well-described competitive phenotype, where *P. aeruginosa* inhibits *S. aureus* growth, ^16^ and the newly described phenotype of coexistence, where *P. aeruginosa* is unable to outcompete *S. aureus* ^19–21^. In order to quantify the importance of this last phenotype, we collected 50 pairs of *S. aureus* and *P. aeruginosa* from 36 co-infected CF patients. The interaction between the two pathogens was analyzed by a competitive test on trypticase soy agar (TSA) plates (Table 1 - fig. 1A). We observed that 61% of strain pairs presented a coexistence phenotype whereas 39% were in competitive interaction. To determine whether the pairs of coexisting strains and competitive strains were phenotypically different, we measured colony size, analyzed the hemolytic properties of each strain, and searched for pigmentation and mucoid phenotype for all *P. aeruginosa* strains. No significant differences were observed between coexisting and competitive strains with respect to pigment production, mucoid phenotype and hemolysis (fig. S1). We observed a significant difference in the size of *S. aureus* colonies in which those of coexisting strains were larger than those of competition strains after 24 h of TSA plate culture. The significance of such a difference and its impact on interaction with *P. aeruginosa* remain to be explored. As others have already described that early infectious strains of *P. aeruginosa* are more aggressive for *S. aureus* than the late infectious strains ^19,20^, we wondered if the type of interaction could be related to the duration of colonization. To answer this question, we determined the duration of co-colonization of *S. aureus* and *P. aeruginosa* for each patient. The average duration of colonization for coexisting strains was 744.8±97.64 days and for competing strains 941.2±137 days. The difference was statistically non-significant (fig. S1).

**Table 1:**
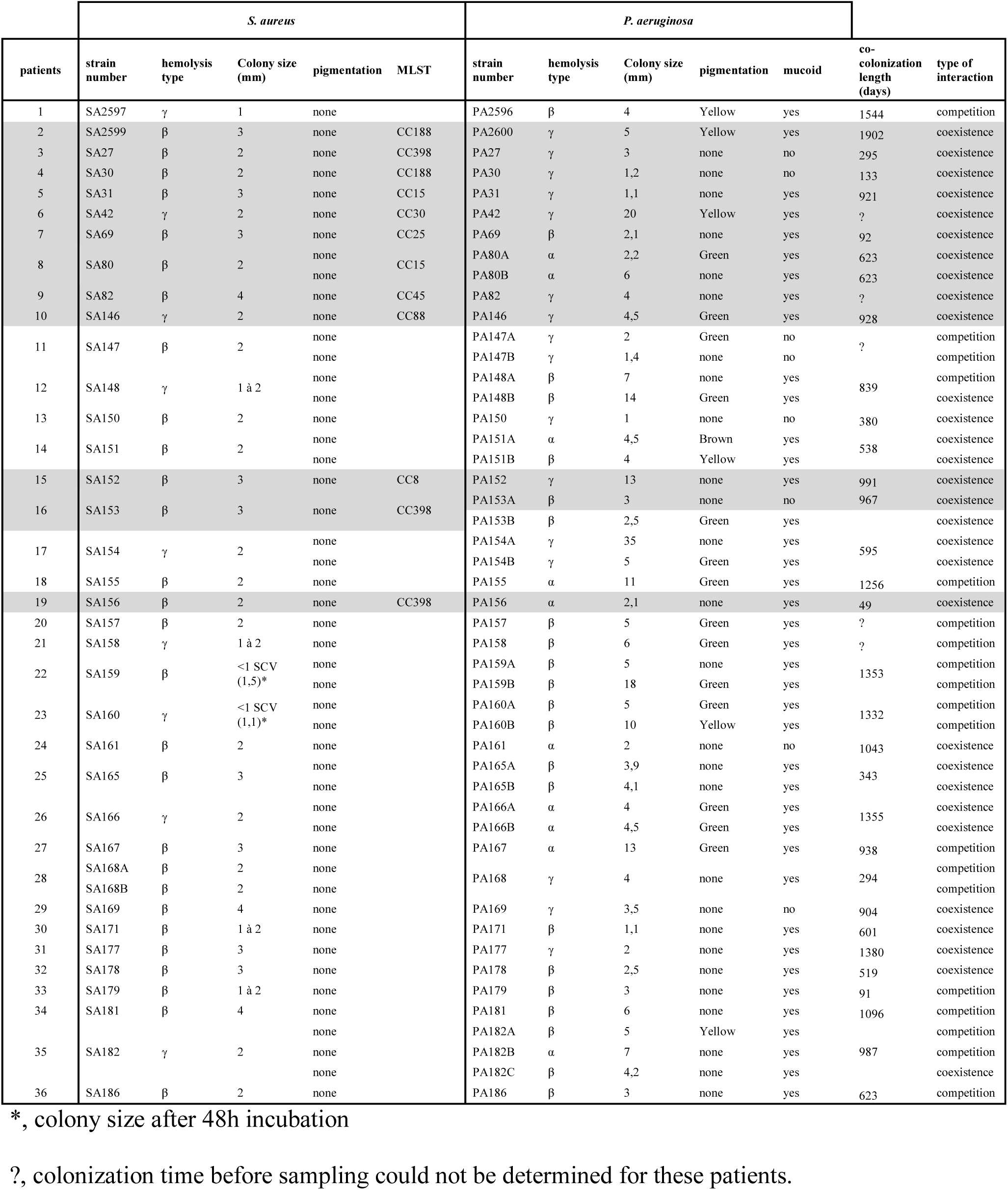
S. aureus and *P. aeruginosa* clinical strains used in this study. 50 couples of *S. aureus* and *P. aeruginosa* were collected from 36 patient sputum samples. Some patients presented several *P. aeruginosa* isolates, and one patient presented two *S. aureus isolates*. Colony size, pigmentation and mucoid phenotype were determined on TSA. Hemolysis type was determined on COS. Interaction type was determined by agar plate competition assay as described in the materials and methods section. The underlined gray isolates correspond to those used for the RNAseq and RT-qPCR analyses. MLST type were determined only for these isolates.

**Figure 1:**
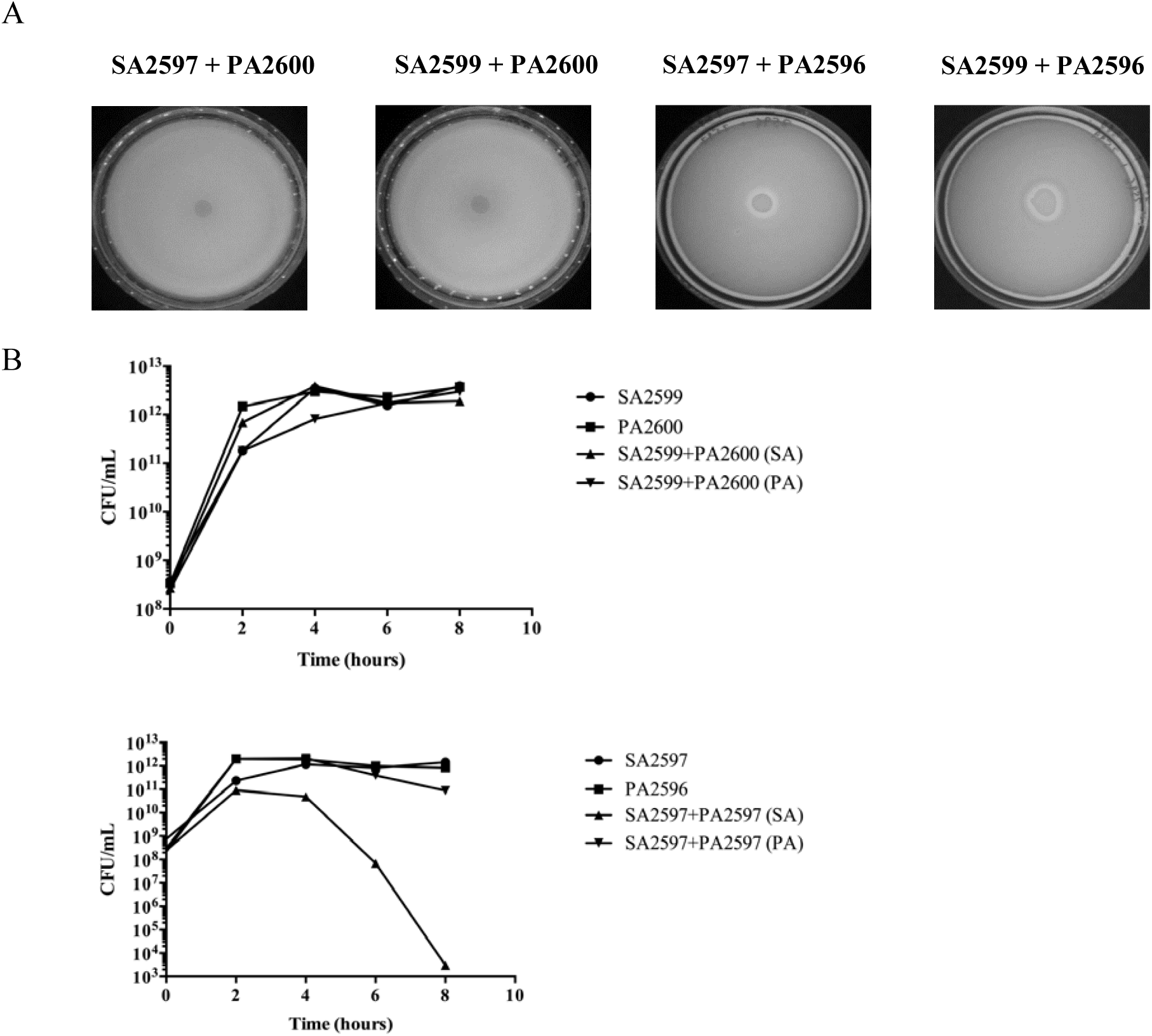
Competition assay between *S. aureus* and *P. aeruginosa*. **A.** Competition test on agar plate. *S. aureus* and *P. aeruginosa* were grown on BHI for 8 hours at 37°C. A layer of *S. aureus* was added on a TSA. After drying, a drop of *P. aeruginosa* was spotted. The inhibition halo indicates a competition state (SA2597+PA2596 and SA2599+PA2596). **B.** Competition assay in planktonic culture. *S. aureus* and *P. aeruginosa* were mono-cultivated and co-cultivated for 8 hours. Every two hours, bacteria were plated on mannitol salt agar (MSA) and cetrimide to count *S. aureus* and *P. aeruginosa*, respectively. The results show one representative experiment from a triplicate. Upper panels, pairs in coexistence. Lower panels, pairs in competition.

Planktonic cultures were conducted on two pairs of strains: one competitive pair (SA2597/PA2596) and one coexisting pair (SA2599/PA2600). In both cases, we observed that *P. aeruginosa* growth was not altered by *S. aureus*. On the other hand, in the case of the competitive pair, *P. aeruginosa* had a negative effect on *S. aureus* growth after 4 hours of coculture (Fig 1B). Agar plate competition assays mixing respectively PA2600 (from coexisting pair) and PA2596 (from competitive pair) with both SA2597 and SA2599 were performed (fig. 1A). PA2596 outcompeted both SA strains whereas PA2600 was unable to inhibit any of the *S. aureus* strains, suggesting that the interaction phenotype is dependent on the *P. aeruginosa* strains. These results were confirmed with other combinations of strains (fig. S2).

### *P. aeruginosa* differentially dysregulates *S. aureus* transcriptome according to coexistence/competition

To obtain an overview of the impact of P. aeruginosa on the expression of *S. aureus* genes, a comparative transcriptomic study was conducted between SA2597 and SA2599 in monocultures, and the same strains in coculture with a competition PA strain (PA2596) and a coexisting PA strain (PA2600). Thus, for each interaction state, we tested two pairs of strains, namely SA2597 / PA2596 and SA2599 / PA2596 for the competition and the SA2597 / PA2600 and SA2599 / PA2600 pairs for coexistence. Gene expression was considered dysregulated when dysregulation was common to both pairs of strains. Therefore, seventy-seven *S. aureus* genes were specifically dysregulated in the context of competition and 140 genes in the context of coexistence while only 16 genes were dysregulated both in competition and in co-existence (Table S4).

KEGG analyses were performed on dysregulated genes to associate each gene with a functional class (fig. 2). In competition state, the main dysregulated class of genes belongs to genetic information and processing, with an increase of tRNA and ribosomal RNA (fig. 2A). We also observed the dysregulation of genes involved in major metabolism pathways of carbohydrates and amino acids. The down-regulation of the Acetyl-coenzyme A synthetase encoding gene *(acsA)* was noted. Other genes involved in energetic metabolism were up-regulated in the presence of *P. aeruginosa*, especially dehydrogenase enzymes such as the lactate dehydrogenase (*ldhA*), the alanine dehydrogenase (*ald1*), the glutamate dehydrogenase (*gluD*), the 1-pyrroline-5-carboxylate dehydrogenase (*rocA*), the 2-oxoglutarate dehydrogenase (*odhA*) and the aldehyde-alcohol dehydrogenase (*adhE*). The upregulation of the *ldh* gene is consistent with the up regulation of the L-lactate permease (*lctP*) encoding gene. All these factors, as well as acetyl-coA, are involved in energetic metabolism and redox reactions conducted to feed the Krebs cycle and ensure the production of ATP.

**Figure 2:**
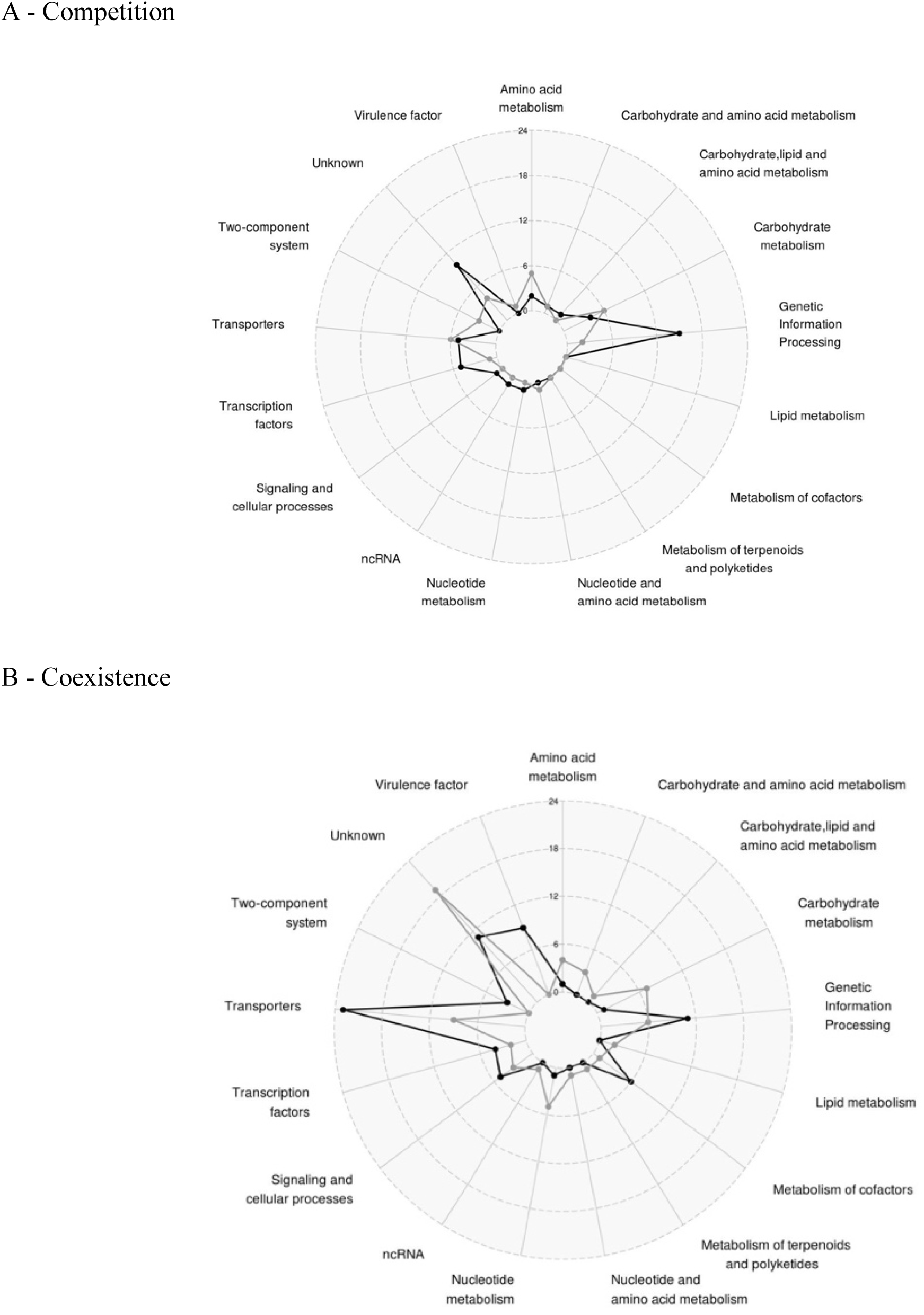
Number and functions of differentially expressed staphylococcal genes in the presence of *P. aeruginosa* when: **A.** both species are in competition (SA2596 and SA2599 were co-cultivated with PA2597) **B.** and in coexistence (SA2596 and SA2599 were co-cultivated with PA2600). RNAs were extracted at 4 hours and a RNAseq was performed in triplicates. KEGG mapper analysis was conducted on common significantly over-expressed (black) and under-expressed genes (grey) to address a functional classification. A gene was considered as differentially expressed when the fold change was strictly higher than 4 with P_adj<0.05.

In the context of coexistence, although *P. aeruginosa* does not appear to impact major metabolic pathways of *S. aureus* as it does not alter growth, the expression of 140 *S. aureus* genes was affected by the presence of *P. aeruginosa* (fig. 2B). Nine known and predicted virulence factor encoding genes were upregulated, including alpha-hemolysin (*hla*), staphylokinase (*sak*), aureolysin (*aur*), the immunoglobulin-binding protein (*sbi*) and staphylococcal complement inhibitor (*scn*) genes. We also observed the overexpression of *saeRS* genes, coding a two component system that has been described as playing a major role in controlling the production of virulence factors such as those mentioned above ^22^.

Other genes whose expression were affected by the presence of *P. aeruginosa* in a coexistence situation are involved in carbohydrate, lipid, nucleotide and amino acid metabolism. Most of them were down-regulated as were several genes (*pgi*, *fbp, fda)* involved in glycolysis and the pentose phosphate pathway. Moreover, two operons (*nrdE*, *nrdI*, *nrdF* and *nrdG*, *nrdD*) belonging to ribonucleotide reductase (RNR) systems and converting nucleoside phosphate into deoxynucleotide phosphate, were both down-regulated (Table S4). RNRs are involved in the *de novo* production of deoxyribonucleotide di- or triphosphates, an essential process for the biosynthesis of DNA and its repair. They catalyze the limiting step of the synthesis of deoxyribonucleotide phosphates and thus control cell concentration ^23^.

Finally, several genes belonging to a transporter family were also over-expressed (polyamines, methionine, iron uptake and antibiotic resistance) in the coexistence state. Notably, all genes from the polyamine operon were over-expressed (*potABCD*) including *potR*, the regulator of polyamine genes. Polyamines control the physiology of *S. aureus* by acting as regulators of several genes involved in metabolism, transport and virulence ^24,25^. In addition, the same pattern was observed for the *metQPN* operon involved in methionine transport and *sirA/B* and *sstA/BC* genes for iron uptake, which may also reflect nutrient competition in coculture. Finally, transporter *norb_3* predicted as belonging to the *nor* family was over-expressed. Pumps from this family export a wide range of antibiotics such as erythromycin, tetracycline and quinolones. Indeed, *norb_3* corresponds to the well-described *tet38* gene involved in tetracycline resistance and internalization in pulmonary epithelial cells ^26,27^.

To confirm these results, we performed co-cultivations with 12 different co-existence *P. aeruginosa-S. aureus* strain pairs from CF patients. The 12 strain pairs came from 12 different patients and presented phenotypic diversity. *S. aureus* isolates belong to 8 different multilocus sequence typing (MLST) types (Table 1). Ten isolates of *P. aeruginosa* were mucoid and four secreted pigmentation, which was representative of the collection of all the isolates. Gene expression was assessed by RT-qPCR for the two categories most impacted: virulence factors and transporters (fig. 3). Regarding virulence factors, we confirmed the over-expression of the aureolysin encoding gene in 6 of the 7 strains that expressed the gene. For *sbi*, 5 out of 12 strains presented over-expression and 5 out 12 presented decreased expression, meaning that there was no clear profile of *P. aeruginosa’s* impact on this gene expression. For the other virulence genes tested, we observed reduced expression in the majority of the strains (10/12 for *hla*, 5/7 for *sak*, 7/12 for *saeRS* and 8/12 for *scn*) (fig. 3).

**Figure 3:**
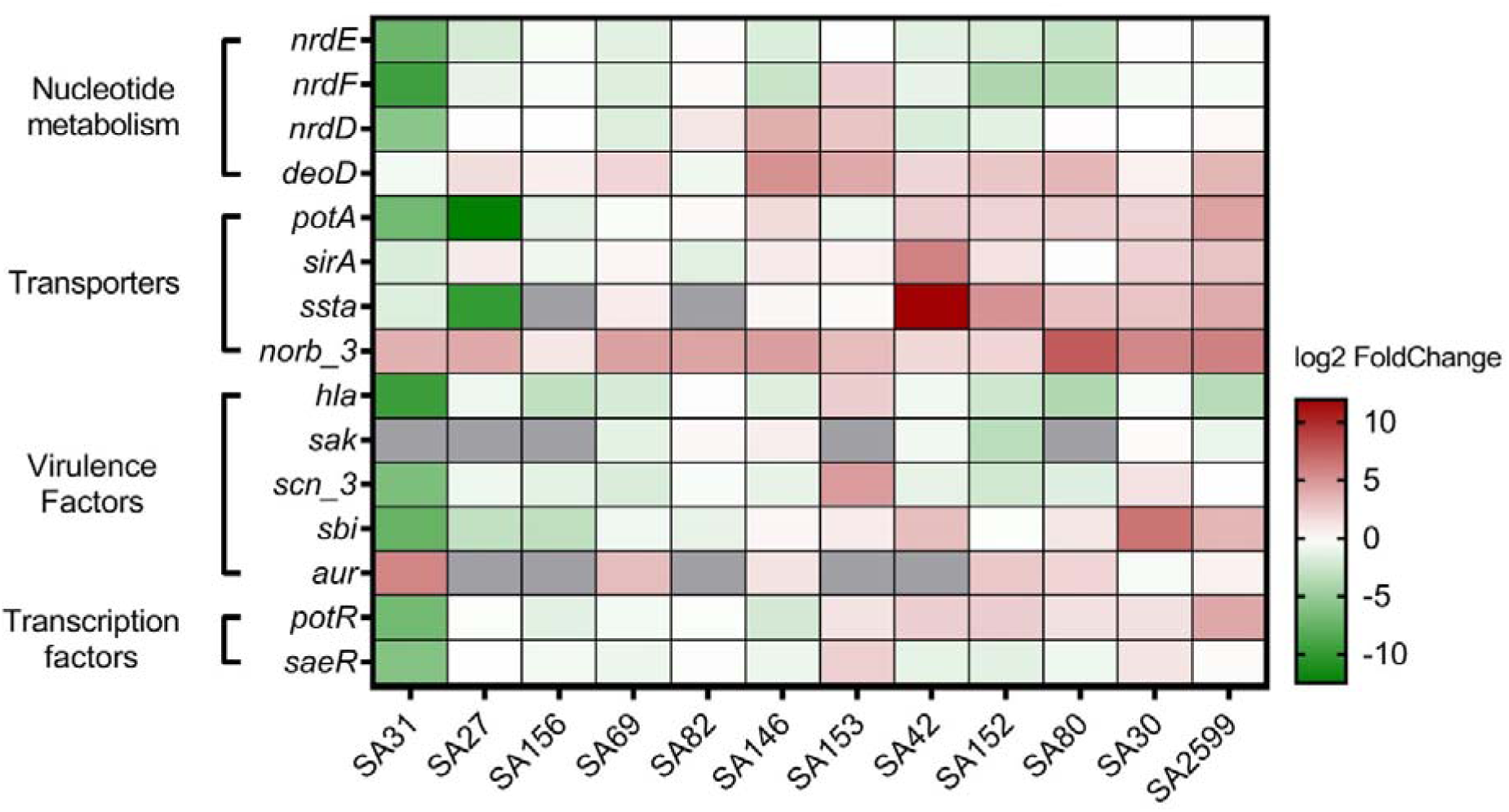
Confirmation of *S. aureus* gene expression dysregulation by *P. aeruginosa*. Twelve clinical SA-PA pairs of strains were co-cultivated for 4 hours. RNAs were extracted and RT-qPCR were performed on 15 genes. The results are represented as fold change of expression (gene relative expression in coculture/ gene relative expression in monoculture) on a heatmap. Under-expressed genes are indicated in green whereas over-expressed genes are indicated in red. No RNA detection is shown in gray. Pairs were hierarchically clustered by the Euclidean method.

For transporter encoding genes, we confirmed the over-expression of *pot* genes and the *sstA* gene in 6/12 and 7/10 strains, respectively. Noticeably, we confirmed the up-regulation of *tet38* genes in 11/12 strains with a fold change ranging from 3 to 200. In addition, *deoD* gene upregulation was also confirmed in 9/12 strains, consistent with its operon structure with *tet38* gene ^26^.

The over-expression of the *tet38* gene is the most predominant transcriptomic alteration in our study and may be of great importance as it can affect the antibiotic susceptibility of *S. aureus*, an important element in the context of CF disease. Therefore, we aimed to better characterize this transcriptomic alteration.

### Over-expression of the *tet38* gene is due to the dysregulation of the MgrA regulatory pathway that impacts other *nor* family genes

To decipher the molecular pathway involved in the over-expression of the *tet38* gene, we analyzed the expression of known regulators in the presence or absence of *P. aeruginosa*. Three transcriptional negative regulators of *tet38* have already been described: TetR21 ^26^, SarZ ^28^ and MgrA ^27^. The expression of these regulators was quantified by RT-qPCR in coculture and compared to expression in monoculture. None of the *tetR21*, *sarZ* and *mgrA* RNA levels was affected by the presence of *P. aeruginosa* (fig. S3).

However, it has been described that regulation by MgrA is dependent on its phosphorylation state ^29^ and that the deletion of *mgrA* induces increased expression of *tet38* ^27^. Therefore, we analyzed the impact of *P. aeruginosa* on *tet38* expression using a Newman Δ*mgrA* mutant (fig. 4). The wild type Newman strain presented a 20-fold change over-expression of the *tet38* gene in the presence of *P. aeruginosa*, as we previously observed in clinical strains. This fold change was reduced to 6 for the Δ*mgrA* mutant. Therefore, it appears that the over-expression of the *tet38* gene is induced by an alteration of the MgrA regulatory pathway. These results were confirmed in *S. aureus* Lac isogenic strains (fig. S4).

**Figure 4:**
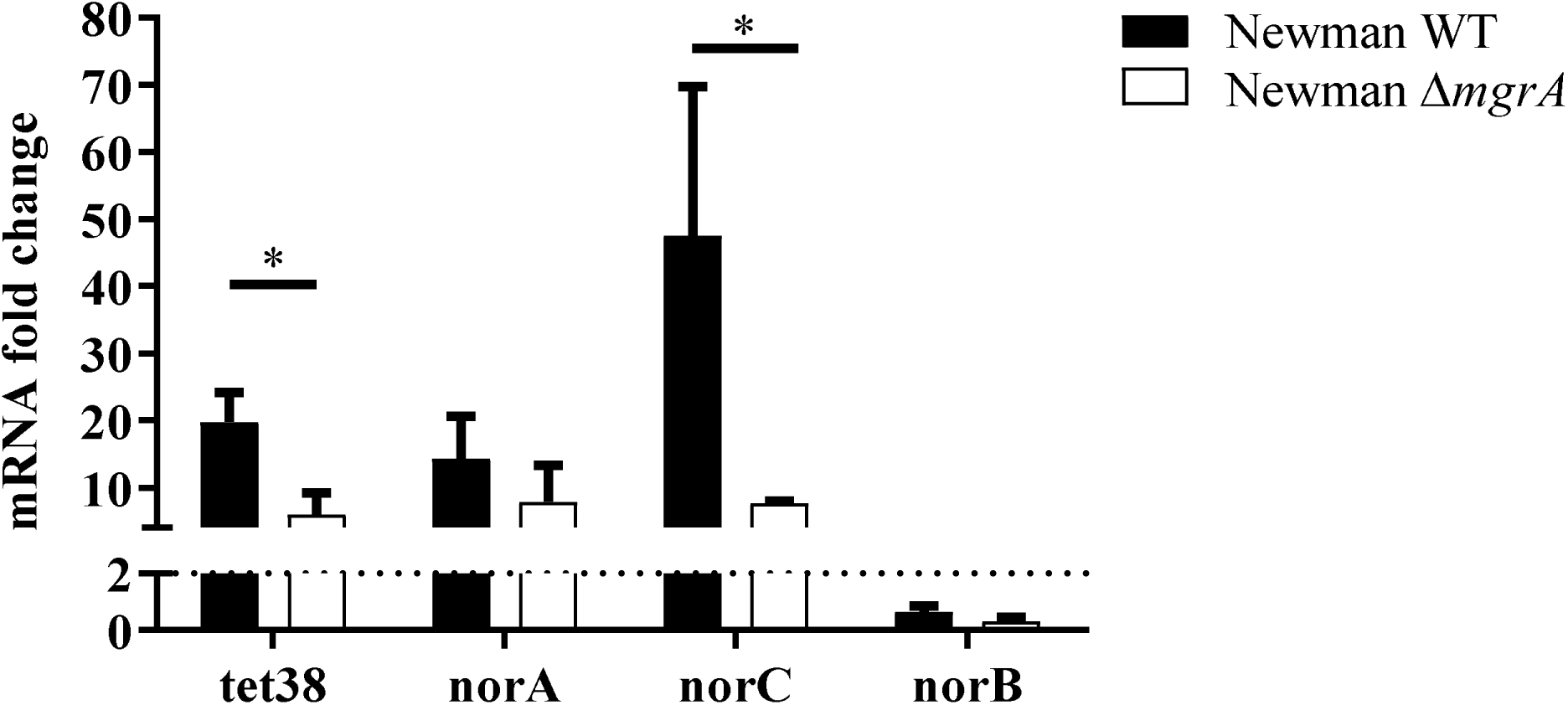
The *S. aureus* MgrA regulator is important for *nor* gene over-expression. Cocultures with *S. aureus* Newman wild type (WT) and Δ*mgrA* mutants and PA30 were performed. RNAs were extracted at 8 hours and *nor* gene expression was monitored by RT-qPCR. The results are shown as the mean + standard deviation of three independent experiments. Dotted lines indicate fold change= 2. Statistical analysis was performed by unpaired t-test (* P<0.05).

MgrA is also a transcriptional regulator of other *nor* family protein genes such as *norA*, *norB* and *norC* ^27,29,30^. Hence, the expression of *nor* genes in *S. aureus* was monitored throughout cocultures of the 12 *S. aureus* strains with *P. aeruginosa* (fig. 5). The *Tet38* gene was significantly overexpressed in the presence of *P. aeruginosa* throughout the 8 hours of the culture. The *NorA* gene was over-expressed in at least 50% of cocultures (6/12 and 7/12) after 4 and 6 hours of culture and *norC* expression was increased at 4 and 8 hours in 6/11 and 7/11 strains, respectively (fig. 5). *norB* was overexpressed only at 8 hours of culture in 50% of the strains tested.

**Figure 5:**
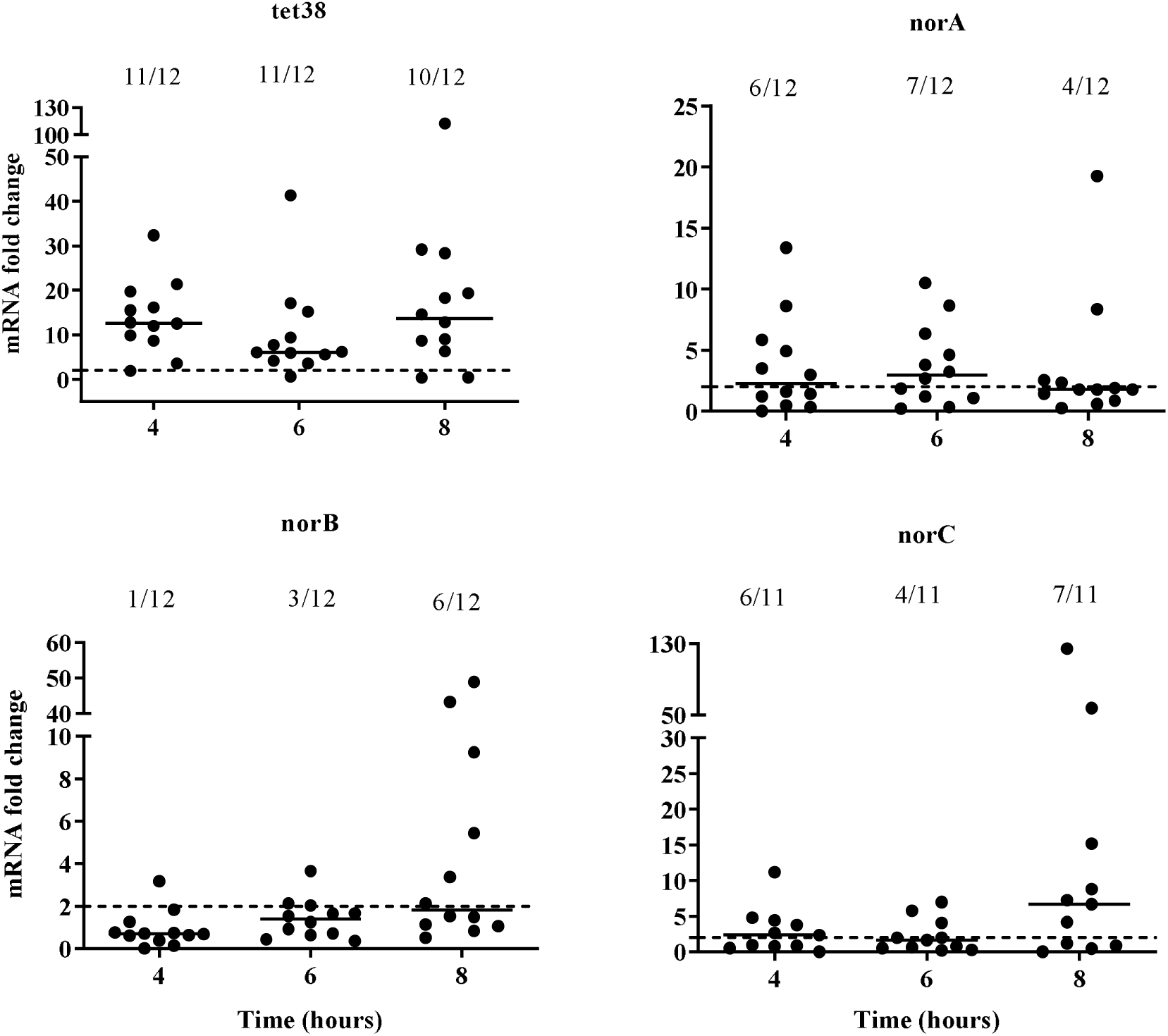
The over-expression of *S. aureus nor* family genes induced by *P. aeruginosa*. Mono- and coculture with twelve clinical strain pairs were performed. RNAs were extracted and gene expression was monitored by RT-qPCR at 4, 6 and 8 hours. The results are represented as fold change expression. Dotted lines indicate fold change= 2. Numbers above each hour indicate the number of pairs with a fold change strictly higher than 2.

From the analysis of expression in the Δ*mgrA* mutant in the *S. aureus* Newman strain, we concluded that over-expression of *norC* was dependent on MgrA integrity (fig. 4). A milder effect was observed on *norA* over-expression. No overexpression of *norB* gene was observed with the *S. aureus* Newman strain. In the *S. aureus* Lac strain, we observed a diminution of *norA* and *norC* overexpression in the Δ*mgrA* mutant but the effect was less significant than on *tet38* gene expression (fig. S4). Therefore, we concluded that the overexpression of *norA* and *norC* genes in the presence of *P. aeruginosa* was partially due to *mgrA* dysregulation.

### The presence of *P. aeruginosa* induces over-expression of *nor* genes by specific and direct interaction

To determine if a secreted product of *P. aeruginosa* induced the over-expression of *nor* genes, a transwell experiment was conducted in which cultures of *P. aeruginosa* and *S. aureus* were separated by a 0.4μm filter. In these conditions, *nor* genes were not overexpressed (fig. 6). The same results were obtained when *S. aureus* culture was exposed to supernatant of *P. aeruginosa* (fig. S3), suggesting that at least a close interaction between the two species was necessary. Finally, to determine if the over-expression was specific to the interaction of *P. aeruginosa*, cocultures were conducted with other bacteria frequently associated with *P. aeruginosa* in CF patients, such as *Burkholderia cepacia* and *Stenotrophomonas maltophilia* (fig. 7). No over-expression was observed, suggesting that the effect was specific to the presence of *P. aeruginosa*. These results were confirmed with other clinical *S. aureus* strains (fig. S6).

**Figure 6:**
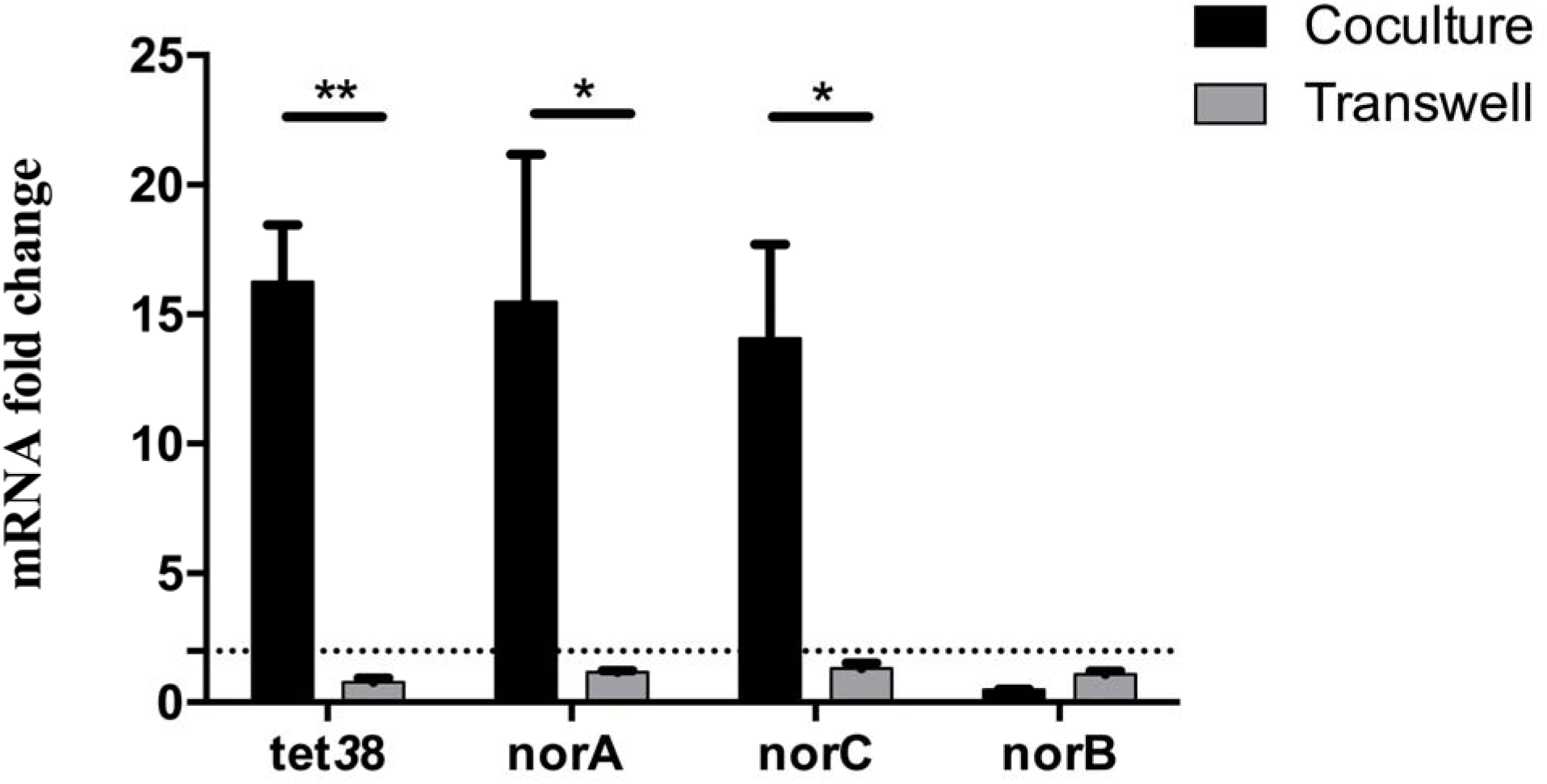
*S. aureus nor* gene over-expression requires close contact with *P. aeruginosa*. *S. aureus* was deposited onto the bottom of wells. *P. aeruginosa* was added either with *S. aureus* (black) or into the insert of transwells (gray). RNAs were extracted and *nor* gene expression was monitored by RT-qPCR. Dotted lines represent fold change= 2. The results are shown as the mean + standard deviation of three independent experiments on SA30-PA30 pairs. Statistical analysis was performed by unpaired t-test (* P<0.05, ** P<0.01).

**Figure 7:**
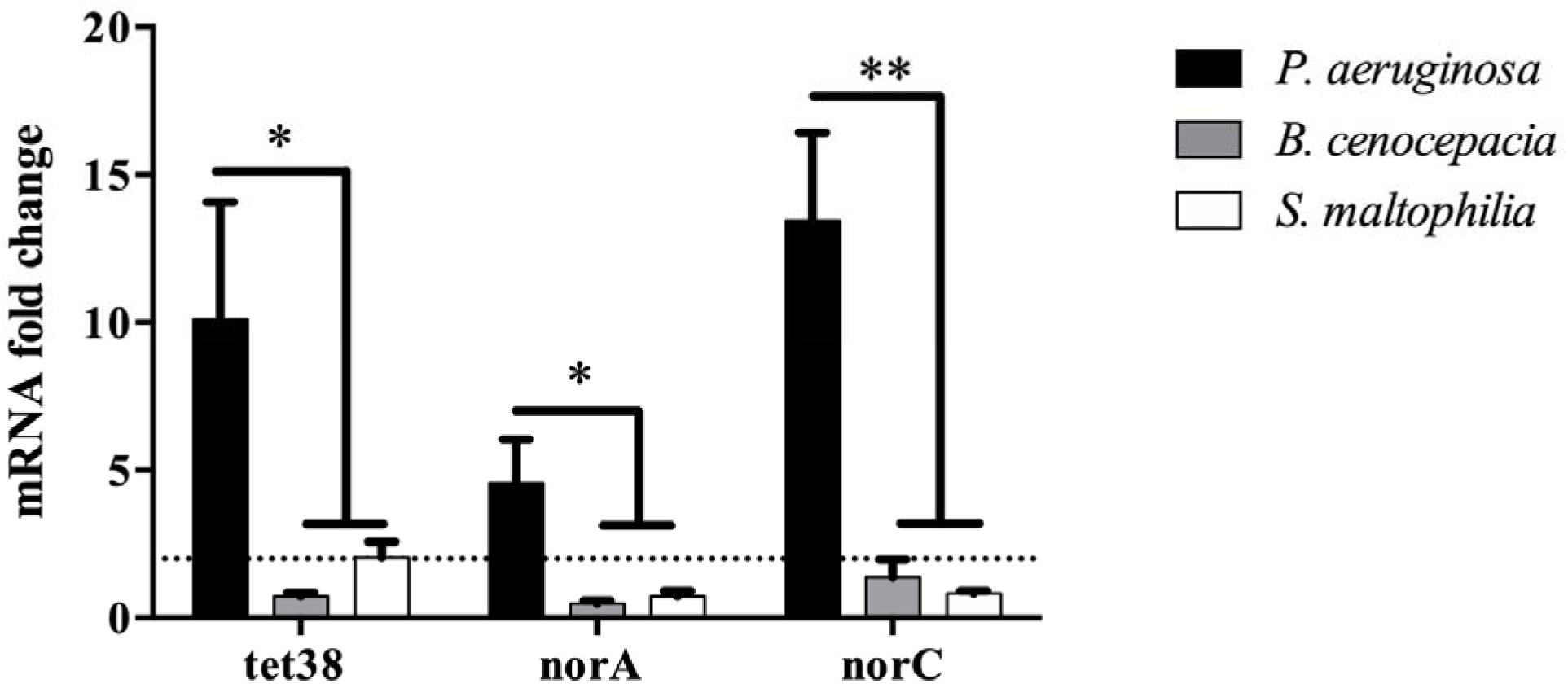
The overexpression of *S. aureus nor* genes is specifically induced by *P. aeruginosa*. *S. aureus* was mono- and cocultivated with *P. aeruginosa, B. cenocepacia* or *S. maltophilia*. RNAs were extracted and gene expression was monitored by RT-qPCR. Dotted lines indicate fold change= 2. The results represent the mean + standard deviation of three independent experiments on SA30-PA30 pairs. Statistical analysis was performed by One-way Anova with Dunnett’s multiple test correction (* P_adj< 0.05, ** P_adj< 0.01).

### Over-expression of *nor* genes induces an increase of antibiotic resistance and internalization of *S. aureus* into epithelial cells

As Tet38 is involved in tetracyclin resistance ^27^ and NorA and NorC are also implicated in quinolones (such as ciprofloxacin) uptake, ^30,31^ the impact of coculture with *P. aeruginosa* on *S. aureus* antibiotic susceptibility was tested. Firstly, the MIC was determined for each of the 12 *S. aureus* strains used (Table S3). Monocultures and cocultures were then exposed to tetracycline and ciprofloxacin at MIC or 2xMIC. After plating on selective agar and numeration, the survival rate was determined by dividing the number of *S. aureus* after antibiotic treatment by the number of *S. aureus* without antibiotic treatment. A 3-fold increase in survival rate was observed at MIC and 2xMIC concentration in the presence of *P. aeruginosa* (fig. 8A and B).

**Figure 8:**
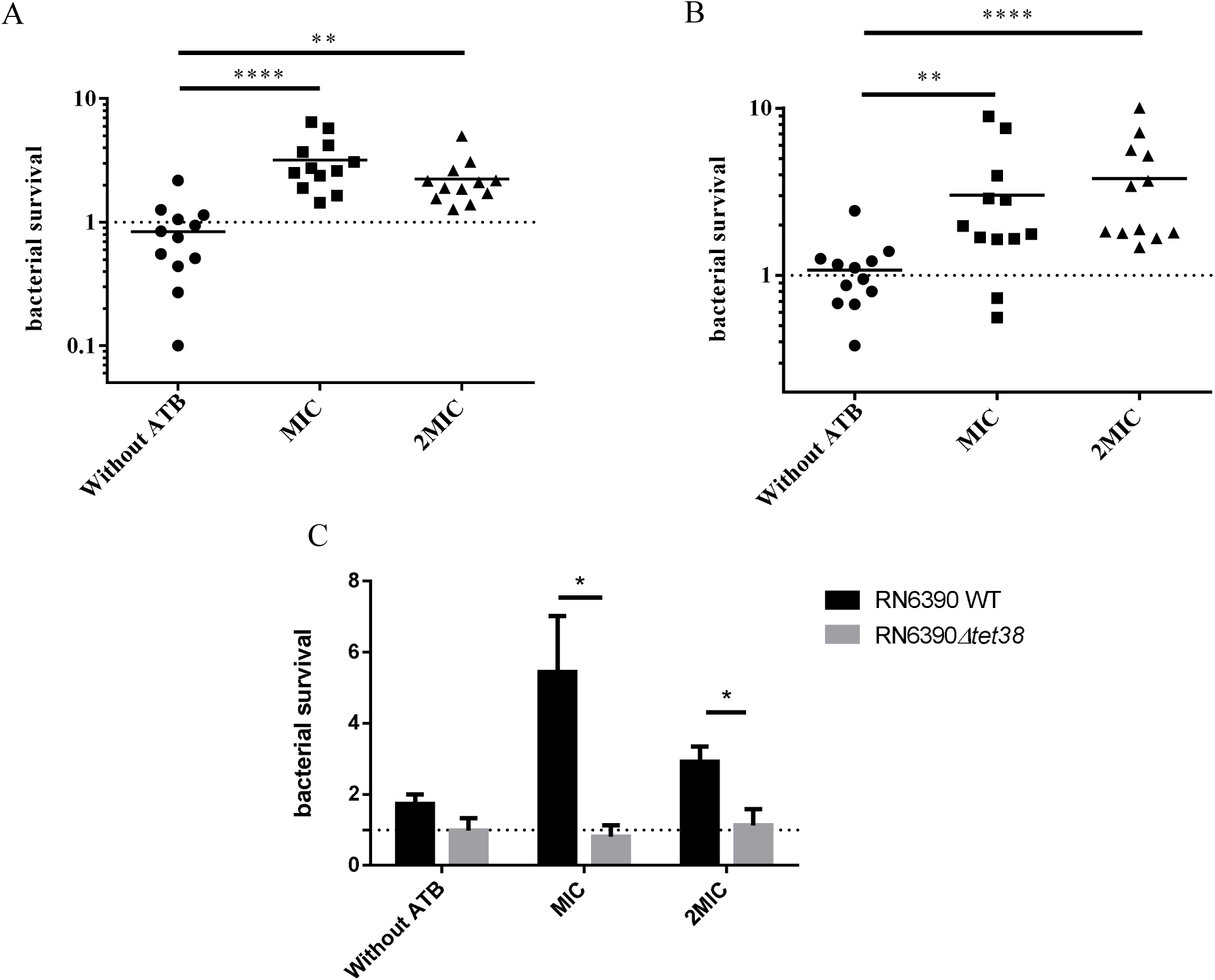
*S. aureus* antibiotic resistance increases when co-cultivated with *P. aeruginosa*. Twelve clinical *S. aureus* strains were mono- and cocultivated with coexisting *P. aeruginosa* strains and exposed to tetracycline (**A**) and ciprofloxacin (**B**) at MIC and 2 x MIC. After 5 hours, cultures were plated on MSA to count remaining *S. aureus*. Bars represent the median and dotted lines bacterial survival equal to 1. Statistical significance was determined by One-way Anova with Dunnett’s multiple test correction (** P_adj<0.01, and **** P_adj<0.0001). **C.** *Tet38* is responsible for the increase of tetracycline resistance induced by *P. aeruginosa*. RN6390 and isogenic Δ*tet38* derivative were cultivated with and without *P. aeruginosa* and susceptibility to tetracycline was monitored. Statistical significance was determined by unpaired t-test (* P<0.05) from three independent experiments. All the results are expressed as the number of surviving bacteria in coculture divided by the number of surviving bacteria in monoculture.

In order to demonstrate that the over-expression of the *tet38* gene was responsible for tetracycline resistance, the impact of *P. aeruginosa* was tested on the RN6390 wild type strain and its isogenic Δ*tet38* mutant upon exposure to tetracycline. As expected, *P. aeruginosa* induced a higher survival rate of the RN6390 wild type strain after tetracycline exposure. On the contrary, it had no impact on the bacterial survival of the Δ*tet38* mutant after tetracycline exposure (fig. 8C), confirming the role of the *tet38* gene in the enhancement of tetracycline resistance in the presence of *P. aeruginosa*.

Tet38 has also been described as being involved in pulmonary epithelial cell internalization ^26^, so the impact of the presence of *P. aeruginosa* on *S. aureus* cell internalization was tested using the Gentamicin protection assay. When A549 epithelial cells were infected with a monoculture of *S. aureus*, no difference was observed in terms of bacterial adhesion onto A549 cells. Five percent of adherent bacteria were internalized into cells. When *S. aureus* was co-cultivated with *P. aeruginosa* before cell infection, 15% of *S. aureus* were internalized, meaning a 3-fold increase of the *S. aureus* internalization rate in the presence of *P. aeruginosa* (fig. 9). To ensure that the effect we observed was not due to an alteration of the A549 cell layer by *P. aeruginosa* that could have facilitated *S. aureus* internalization, we performed an LDH measurement on the cell supernatant as an indicator of A549 cell viability (fig. S7B). Although the LDH level was slightly higher for cells infected only with *P. aeruginosa*, we found no significant difference between the *S. aureus* infected and *S. aureus* plus *P. aeruginosa* co-infected cells. Indeed, the A549 co-infected cells had the lowest level of LDH. Moreover, microscopic observation of the cells revealed no difference between the mono- and co-infected cells (fig. S7A). Therefore, it appeared that the presence of *P. aeruginosa* did not alter the A549 cells and the highest rate of *S. aureus* internalization was due to its direct impact on *S. aureus*. However, we could not be sure that the increase in the internalization rate was directly related to *tet38* overexpression. It could be the result of a modification of different factors involved in the internalization process, although we did not identify such factors in our transcriptomic analysis apart from the *tet38* gene.

**Figure 9:**
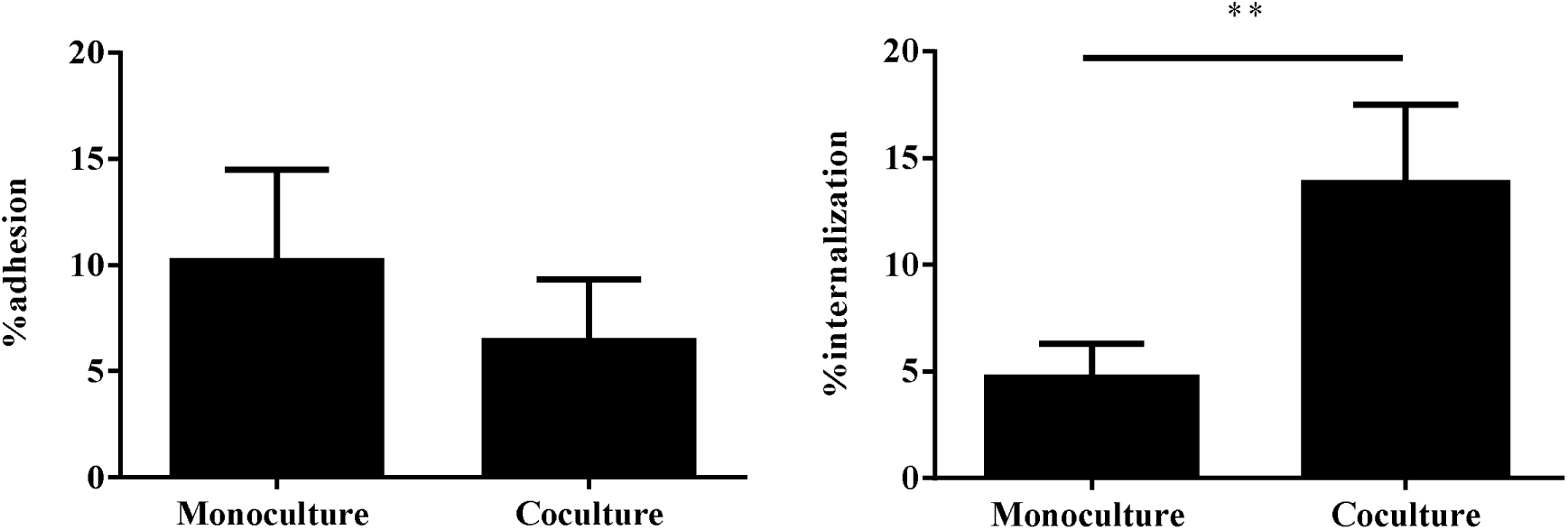
*S. aureus* internalization is increased within A549 epithelial pulmonary cells in the presence of *P. aeruginosa*. **A.** Adhesion of *S. aureus* onto epithelial cells. A549 cells were infected at MOI 10:1 for *S. aureus* monoculture and 20:1 for *S. aureus*/*P. aeruginosa* coculture. After 2 hours of contact, cells were washed with phosphate buffer saline (PBS) to remove unattached bacteria and lysed with sterile water. Supernatants were plated on MSA to count *S. aureus*. The results are represented as the percentage of inoculum that adhered. **B.** Internalization of *S. aureus* within epithelial cells. After 2 hours of contact, cells were treated with antibiotics and lysostaphine for one hour, lysed with sterile water and bacteria plated on MSA. The results are represented as the percentage of adhered cells that have internalized. All values represent the mean + standard deviation from three independent experiments with three strain pairs (SA27-PA27, SA31-PA31 and SA69-PA69). Statistical significance was determined by unpaired t-test (** P<0.01).

## DISCUSSION

The goal of this study was to investigate the impact of the interaction of coexisting *S. aureus* and *P. aeruginosa* on *S. aureus* at the transcriptional and phenotypical levels.

Firstly, we collected isolates from co-colonized CF patients and demonstrated that in 61% of cases, *S. aureus* was able to coexist with *P. aeruginosa* with no alteration of its growth. So far, it appears that coexistence of the two pathogens may be a frequent situation in the context of CF patients’ lung infection. Previous studies described that early infectious strains of *P. aeruginosa* are more aggressive for *S. aureus* than the late infectious strains ^19,20^. In the present study, we did not find any correlation between the interaction type and the duration of *S. aureus* and *P. aeruginosa* co-colonization. In the first studies, limited numbers of patients were studied (1 in Michelsen et al., 2014 and 8 in Baldan et al., 2014). Even in our present study, only 11 patients had competitive strain pairs, which might be not be enough to reach a conclusion. Up to now, it has been difficult to conclude whether the interaction type between the two species is linked to the evolution of the *P. aeruginosa* strain over the time of co-colonization. A larger cohort of patients would be needed to answer this question. Also, longitudinal clinical studies would be appropriate to analyze the kinetics of interaction evolution over time and determine how it could affect patients’ health. Furthermore, the conditions and environmental factors leading to co-existence instead of competition require clarification, particularly through studies such one conducted recently that demonstrated the positive impact of hypoxia found in static mucus within CF airways on a coexisting interaction ^32^.

The type of interaction may have an impact on the physiology of the two pathogens involved. In order to answer this question, we conducted a transcriptional study of the impact of *P. aeruginosa* on *S. aureus*.

In the context where *P. aeruginosa* inhibits *S. aureus* growth, transcriptomic modifications affect major metabolism pathways such as translation, Krebs cycle and genes involved in oxidative stress. The increase in the amount of tRNAs and ribosomal RNAs observed could be attributed to a decrease in translation efficiency. Regarding energetic metabolic pathways, we observed a down-expression of Acetyl-coA synthetase, a key factor metabolized into pyruvate to feed the Krebs cycle and produce energy. The down-regulation of expression observed may certainly lead to a defect in ATP production. Conversely, we observed the increased expression of several dehydrogenase enzymes, suggesting a shift from aerobic respiration to lactic acid fermentation to feed the Krebs cycle, as shown previously in laboratory strains ^33,34^. Certain dehydrogenases, such as *adhE* and *gluD* genes, are also implicated in oxidative stress responses ^35^. All these major dysregulations observed are consistent with the lethal effect of *P. aeruginosa* on *S. aureus* in competitive interaction.

More genes were dysregulated when *S. aureus* and *P. aeruginosa* were coexisting. We observed a drastic modification in the nucleotide synthesis pathway with a down-regulation of genes involved in the *de novo* pathway (*nrd* operon) and upregulation of the *deoD* gene encoding a purine nucleoside phosphorylase involved in an alternative metabolic pathway for nucleotides when the *de novo* pathway is altered. We also observed a down-expression of genes involved in the classical energetic metabolism pathways: glycolysis and pentose phosphate pathways (*pgi, fbp, fda*). These results suggest nutritional competition between the two pathogens and indicate that in our conditions, *S. aureus* preferentially produced energy and nucleotides from sources other than glucose.

Finally, we observed the increased expression of several transporters, especially *tet38*, *norA* and *norC* genes. Curiously, the *tet38* gene belongs to the same transcription unit as the *deoD* gene. It is tempting to speculate that the overexpression of *tet38-deoD* operon may be linked to the down regulation of the *nrd* genes to compensate for the alteration of the *de novo* nucleotide synthesis pathway.

These genes are members of the Nor family and encode efflux pumps involved in antibiotic resistance. *Tet38* was the most impacted gene with 11 pairs of 12 for which we observed an increased expression throughout the coculture kinetics, whereas the over-expression of other *norA* and *norC* genes appeared on 7 pairs of 12 and 11of 12 at 6 and 8 hours, respectively. Given that the pair 2599/2600 used for the RNA sequencing (RNAseq) was unable to upregulate *norA* and *norC* at 4 hours of coculture, it was expected that *norA* and *norC* genes would not appear in the RNAseq results. The over-expression of *tet38*, *norA* and *norC* genes appeared to be at least due to a dysregulation of the MgrA pathway. Indeed, the Δ*mgrA* mutant provoked a strong effect on *tet38* over-expression but only a slight effect on *norA* and *norC*. Thus, *mgrA* seems to be essential for *tet38* over-expression and other regulators must be implicated for the *norA and norC* genes. In addition, we were unable to observe clear *norB* up-regulation in the presence of *P. aeruginosa*. Indeed, MgrA act as a repressor of *tet38*, *norA* and *norC* and an activator of *norB* in an *rsbU* positive background strain ^36,37^. This discrepancy may explain our results. Despite its role in *tet38* induction during coculture, *mgrA* expression was not affected by the presence of *P. aeruginosa*. However, it has been shown that the phosphorylation state of MgrA, regulated by RsbU and PknB factors, was a key mechanism for regulation of *nor* family gene expression ^29^. Thus, *P. aeruginosa* may induce a variation of MgrA phosphorylation leading to a modification of *nor* gene expression.

*Nor* proteins are responsible for antibiotic efflux (tetracycline and fluoroquinolone), and we demonstrated that *P. aeruginosa* increased the survival rate of *S. aureus* after exposure to tetracycline and ciprofloxacin. For tetracycline, the effect appears to be mainly due to *tet38*, as antibiotic resistance in presence of *P. aeruginosa* was eliminated in a *tet38* mutant. The same analysis could not be performed for *nor* genes due to the functional redundancy of the *norA, norB* and *norC* genes and the difficulty in obtaining triple mutants. Tet38 is also able to interact with the CD36 receptor on pulmonary epithelial cells to favor *S. aureus* internalization ^38^. Indeed, in the presence of *P. aeruginosa*, we observed a higher rate of *S. aureus* internalization into epithelial cells. Internalized bacteria are more resistant to antibiotics and less detectable by the immune system ^39^. Our results suggest that by coexisting with *P. aeruginosa*, *S. aureus* could hide from the host immune system and be more resistant to antibiotics.

We did not identify the *P. aeruginosa* specific signal responsible for *S. aureus* gene expression dysregulation. However, we demonstrated that it seems to be specific to *P. aeruginosa* (no other species tested had the same effect) and requires very close proximity between *S. aureus* and *P. aeruginosa* to be effective. Transcriptomic analysis revealed that the *S. aureus potRABCD* operon for polyamine uptake and regulation exhibited significant fold change upon exposure to *P. aeruginosa*. The same effects were observed by Yoa and Lu ^25^ when exposing *S. aureus* to polyamines. Moreover, the exposure of *S. aureus* to spermine induces transcriptional modifications including over-expression of antibiotic efflux pumps such as *norA* and *tetM* genes and the decreased expression of many genes involved in carbohydrate metabolism and transport^24^. These results are consistent with the reduced expression of genes involved in glycolysis and the pentose phosphate cycle described previously ^24,25^. Indeed, we observed the same profile after exposure to *P. aeruginosa* as other authors observed after exposure to spermine. Finally, *P. aeruginosa* presents polyamines at the outer surface of its membrane, more precisely putrescine and spermidine ^40^. Therefore, we suggest that the *P. aeruginosa* polyamines present at the outer surface may be a signal for *S. aureus* transcriptional modifications. Further investigation will be necessary to confirm this hypothesis.

To the best of our knowledge, this study is the first to characterize the transcriptomic profile of coexisting *S. aureus* and *P. aeruginosa* pairs in a clinical context. We demonstrate that this commensal-like interaction induces phenotypical changes in *S. aureus* such as increased antibiotic resistance and host cell internalization. These phenotypes may favor the persistence of *S. aureus* in the context of chronic infection. Since this state of coexistence is apparently solely attributable to *P. aeruginosa*, the selective advantage for *P. aeruginosa* leads to questions. Indeed, previous studies showed that cocultivation with *S. aureus* induces LPS mutation in *P. aeruginosa* associated with fitness gain and antibiotic resistance ^41^, and that *S. aureus* exoproducts restore and enhance *P. aeruginosa* motility ^32^. The state of coexistence could thus represent a trade-off allowing both pathogens to benefit mutually and maintain equilibrium. However, the impact of *S. aureus* on *P. aeruginosa* in this state of coexistence warrants further investigation.

## MATERIALS AND METHODS

### Bacterial strains and culture growth

The bacterial strains and plasmids used in this study are listed in Tables 1 and S1. The clinical strains were originally isolated by the Institute for Infectious Agents from sputum samples of patients followed-up in the two CF Centers of Lyon (Hospices civils de Lyon, France). The strains were collected between May 2016 and June 2017 from 36 different patients. The size of the colonies, pigmentation and mucoid phenotype were determined after 24h of culture on TSA. Hemolysis type was determined after 24h of culture on Columbia agar (COS). MLST clonal complex assignment was inferred from microarray analysis ^42^.

The Δ*tet38* mutant of *S. aureus* RN6390 strain was obtained using the pMAD vector ^43^. The two DNA fragments corresponding to the chromosomal regions upstream and downstream of the *tet38* coding sequence were amplified by PCR using primers listed in Table S2. They were subsequently cloned into the pMAD vector using the In-Fusion® HD Cloning Kit (Clonetech). The resulting plasmid was electroporated into the RN4220 recipient strain and then transferred to RN6390. Growth at non-permissive temperature (44°C) was followed by several subcultures at 30°C and 37°C to promote double crossing over as previously described ^44^.

All the strains were grown in Brain Heart Infusion (BHI, BBL™ Difco) with shaking at 200 rpm at 37°C overnight. Cultures were diluted to 0.1 OD_600nm_ and incubated for 2.5 hours (37°C, 200rpm). Bacteria were spun down at 4000 rpm for 10 min and re-suspended in fresh BHI medium to 2 OD_600nm_. Ten ml of *S. aureus*, *P. aeruginosa*, *B. cenocepacia* and *S. maltophilia* suspension were added to 10 ml of BHI for monocultures. Ten ml of *S. aureus* suspension was mixed with respectively 10 ml of *P. aeruginosa*, *B. cenocepacia* or *S. maltophilia* for cocultures. Cultures were grown for 8h. Every two hours, cultures were plated on mannitol salt agar (MSA, BBL™ Difco) and cetrimide (Difco™) for *S. aureus* or *P. aeruginosa* counts, respectively. For supernatant exposure, 10 mL of *S. aureus* culture was added to 10 ml of supernatant from 8 hours culture of *P. aeruginosa*.

Transwell^®^ (Corning) preliminary experiment demonstrated that bacteria were not able to cross the 0.4 μm filter of the insert. *S. aureus* and *P. aeruginosa* suspensions from 2.5 h culture were pelleted and re-suspended to OD_600nm_ =1 for *P. aeruginosa* and OD_600nm_ =0.33-0.5 for *S. aureus*. The Transwell^®^ experiment was carried out as previously described ^45,46^ with a few modifications. For wells without insert, 400 µL of *S. aureus* OD_600nm_ =0.5 suspension and 200µL of either BHI or *P. aeruginosa* were added. For wells with insert, 600 µL of *S. aureus* 0.33 OD_600nm_was deposed into the wells while 200 µL of either BHI or *P. aeruginosa* was placed onto the insert. The Transwell^®^ system was incubated at 37°C for 8 hours.

### *Staphylococcus aureus* growth inhibition on TSA

From overnight cultures, *S. aureus* and *P. aeruginosa* suspensions were diluted to OD_600_ =0.5 and 100 µl of *S. aureus* suspension was spread uniformly onto TSA plates. Then, 5 µl of *P. aeruginosa* was added at the center of the plates. The plates were incubated at 37°C. The competitive phenotype was characterized by an inhibition halo of *S. aureus* growth, which was measured. The strains were considered as coexisting in the absence of inhibition halo.

### Genome sequencing and annotation

Sequencing libraries were prepared from 1 ng of SA2597 and SA2599 DNA extracted using the DNA Isolation Kit (MO BIO). Library preparation was performed with the Nextera XT DNA sample preparation kit (Illumina) and index kit (Illumina). Library validation was performed on a 2100 Bioanalyzer (Agilent Technologies) to control the distribution of fragmented DNA. WGS was performed with an Illumina HiSeq (Illumina) to generate 150-bp paired end reads. Genomes were sequenced with an average coverage of 130x. Adapters and other illumina-specific sequences were cut from the reads for each set of raw data. Furthermore, Trimmomatic v0.36^47^ was used to perform an additional trimming step using a sliding window with an average quality threshold of 20. Data were checked for quality by FastQC v0.11.6 (S. Andrews, 2010. FastQC: a quality control tool for high throughput sequence data; available online at: http://www.bioinformatics.babraham.ac.uk/projects/fastqc). Assemblies were performed using SPAdes v3.11.1 ^48^. Contigs smaller than 200 bp or with a coverage threshold smaller than 2 were removed manually. Assembly quality control was performed using Quast v4.6.1 ^49^. Genome annotation was processed through Prokka v1.13 including ncRNA prediction ^50^.

To compare CDS and ncRNA from SA2597 and SA2599, the N315 strain (NC_002745.2) was used as a reference. Refseq numbers were gathered from N315 and used as ID tags for common genes. For non-common genes, CDS and ncRNA from SA2597 and SA2599 were blasted with each other with a coverage and identity of 90%. Finally, refseq numbers were also used to gather KEGG numbers and perform functional classification with the Kyoto database. The complete genome sequences for the SA2597 and SA2599 strains were deposited in GenBank under the accession numbers GCA_005280135.1 and GCA_005280145.1.

### Transcriptomic analysis

Cultures and transcriptome sequencing were performed in duplicates or triplicates. The OD_600_ of each culture was normalized to 1.0 at a time of 4 hours for the mono and cocultures. One mL was centrifuged for 5 minutes at 13,000 rpm. Bacteria were treated with lysostaphin (2.5 mg/mL) and lysozyme (50 mg/mL) prior to RNA extraction using the RNeasy Plus Mini Kit (Qiagen). RNAs were treated with TURBO DNA-*free*™ (Invitrogen). rRNAs were depleted using the Ribo-Zero rRNA Removal Kit (Illumina). The cDNA libraries were compiled using the TruSeq Stranded Total RNA Library Preparation Kit (Illumina). The quantification and quality of the DNA libraries was evaluated by Bioanalyzer. The libraries were sequenced using Illumina Hi-Seq 2500 with High-Output (HO) mode, using a V4 chemistry sequencing kit (Illumina). Reads were then processed to remove adapter sequences. Poor quality reads were excluded by Trimmomatic ^47^, using a sliding window with an average quality threshold of 20. Each RNAseq read sample was mapped against its own genome through Bowtie2 v2.3.0 with a sensitive local alignment method^51^. Output files were sorted by read names and converted into BAM format using Samtools v1.3.1. Reads were counted on all feature types (CDS, nc/t/tm/rRNA) using a union mode on Htseq-count v0.6.1 software ^52^. To estimate the enrichment values for the differential expression analysis, statistical analysis was done using R v3.3.3 and DEseq2 v1.14.1 ^53^. Gene expression was considered as dysregulated when: (i) the fold change between co-culture and monoculture was at least 4-fold, (ii) the dysregulation was observed in the two pairs of strains, (iii) the dysregulation was specific to coexistence or competition state. The RNAseq data that support our findings are available in the SRAdatabase under the BioprojectID PRJNA552713, PRJNA552715, PRJNA552786, PRJNA554237, PRJNA554233, PRJNA554237.

### RT-qPCR

RNA extractions were performed using the RNeasy Plus Mini Kit (Qiagen). A DNAse treatment was performed on 10 μg of RNAs using TURBO DNA-*free*™ (Invitrogen). The absence of contaminating gDNA was controlled by PCR. cDNA was synthetized from 1µg RNA using the Reverse Transcription system kit (Promega). The qPCR reactions were performed with PowerUp™ SYBR™ Green Master Mix (Thermofisher) following the manufacturer’s instructions. The target genes and primers used are listed in Table S2. The housekeeping genes *gyrB* and *hu* were used as endogenous control. Gene expression analyses were performed using the ΔCt method.

### Antibiotic resistance assay

MICs of tetracycline and ciprofloxacin (Sigma) were determined by BHI micro-dilution (Table S3). For the antibiotic resistance assay, 4 hour mono-cultures of *S. aureus* and cocultures of *S. aureus*/*P. aeruginosa* were diluted to OD_600nm_ =0.002 or 0.004, respectively, and exposed to antibiotics at MIC and 2xMIC for 5 hours at 37°C at 200 rpm in 1mL of BHI. Cultures were plated on MSA agar plates for *S. aureus* counts. The percentage of bacterial survival after antibiotic treatment was determined by dividing the number of *S. aureus* after antibiotic treatment by the number of *S. aureus* without antibiotic treatment.

### Internalization within A549 cells

*S. aureus* monocultures and *S. aureus/P. aeruginosa* cocultures were performed for 4 hours as previously described ^54^. A549 cells were grown in DMEM GlutaMAX™ medium (Gibco) supplemented with 10% of Fetal Bovine Serum (37°C, 5% CO_2_). 24-well tissue culture plates were seeded at 80 000 cells per well. After 24 hours, the cells were washed twice with 1 ml of Phosphate Buffered Saline (PBS, Gibco) and infected at a multiplicity of infection (MOI) of 10:1 for mono-culture and 20:1 for coculture. Cells were incubated for 2 hours, then washed once in PBS to remove non-adherent bacteria and incubated for 1 hour in DMEM GlutaMAX™ supplemented with 400 µg/mL gentamicin (Sigma), 100 µg/mL polymyxin B (Sigma) and 10 µg/mL lysostaphin to kill extra-cellular bacteria. Cells were washed again with PBS once and lysed with deionized water. Cell lysates were plated on MSA to quantify the intracellular bacteria.

### Data availability

The datasets generated during and/or analyzed during the current study are available from the corresponding author on reasonable request.

The complete genome sequences for the SA2597 and SA2599 strains have been deposited in GenBank under the accession number GCA_005280135.1 and GCA_005280145.1.

The RNAseq data that support our findings are available in the SRAdatabase under the BioprojectID PRJNA552713, PRJNA552715, PRJNA552786, PRJNA554237, PRJNA554233, PRJNA554237.

### Ethical statement

All the methods were carried out in accordance with relevant guidelines and regulations.

## Acknowledgements

This work was supported by the Fondation pour la Recherche Médicale, grant number ECO20170637499 to LC; Finovi foundation to KM and FV. We thank Xavier Charpentier of CIRI, Lyon-France, for discussing the work and providing valuable advice.

## Author contributions statements

PB, LC, SB, FV and KM designed and analyzed the experiments. PB conducted the experiments. SB conducted and analyzed all bioinformatics works. ADJ collected the clinical samples from CF patients. KM and FV coordinated the project. PB and KM collected the data and wrote the first draft of the manuscript. All the authors contributed to manuscript revision and read and approved the submitted version.

