## Supplementary_data for "Coexistence with *Pseudomonas aeruginosa* alters *Staphylococcus aureus* transcriptome, antibiotic resistance and internalization into epithelial cells"

**Table S1:** Laboratory *S. aureus* strains, other species strains and plasmids used in this study.

| **Strains and Plasmids** | | **Genotypes** | **References** |
|---|---|---|---|
| *S. aureus* | LUG3040 | RN6390 WT | [55] |
| | LUG2955 | RN6390 Δ*tet38* | This study |
| | LUG26 | Newman WT | [56] |
| | LUG1790 | Newman *RNAIII*::*tetM* | [57] |
| | LUG30 | Newman *agr*::*tetM* | [58] |
| | LUG3135 | Newman *mgrA*::*tetM* | [59] |
| | LUG2262 | Lac WT | [60] |
| | LUG3134 | Lac *ΔmgrA* | [59] |
| | LUG2995 | SF8300 WT | [61] |
| | LUG2987 | SF8300 *ΔagrA* | [61] |
| *B. cepacia* | LUG2886 | CF clinical strains | This study |
| *S. maltophilia* | LUG2884 | CF clinical strains | This study |
| Plasmids | pMAD | pMAD vector | [43] |
| | pMAD-*tet38* | pMAD for *tet38* allelic exchange | This study |

**Table S2: List of primers used in this study**

| Primers | Direction | Sequences (5'-3') | Target | Size (bp) |
|---|---|---|---|---|
| OLUG4-F | Forward | AGCGAGTGAAAACACGCAAC | *sbi* | 152 |
| OLUG4-R | Reverse | TTCTTGTGCACGTTCTGGGT | | |
| OPB1bF | Forward | CAAACTGCCGGTTTAGCTGC | *norb_3* | 103 |
| OPB1bR | Reverse | TGCTCGTACTTAAGCCAAGGT | | |
| OPB2F | Forward | ACAATTTGGCACACCAACAGA | *potA* | 100 |
| OPB2R | Reverse | TCTAACCATGCGTCCTTCAACA | | |
| OPB5F | Forward | ACGCAAGAAGAACTTGCTGAAC | *potR* | 124 |
| OPB5R | Reverse | GCGTCGTTCCTAACACCTCT | | |
| OPB19bF | Reverse | CAACACGTTGCCATTGTTGC | *nrdE* | 149 |
| OPB19bR | Forward | GACACTAGCTCACCACGACG | | |
| OPB24F | Reverse | TCCGCCAGCTAAGTTCCAAG | *sirA* | 142 |
| OPB24R | Forward | TCCAAATGCGAAAGATGCTGC | | |
| OPB31F | Reverse | AACGTTGGATGCGTATGGGT | *deoD* | 88 |
| OPB31R | Forward | AATGCTTCGACACCAGCGTA | | |
| OPB32F | Reverse | AGCTGAAGCGACTTTGTCAGATGC | *mgrA* | 110 |
| OPB32R | Forward | AGCGTGAACGTTCCGAAGTCGA | | |

| | | | | |
|---|---|---|---|---|
| SarZ-F1<br>SarZ-R1 | Forward<br>Reverse | GATTCTGGAACACTGACACCAT<br>AGCAAGAGGGCTTTTATTGCT | *sarZ* | 138 |
| OPB35F<br>OPB35R | Forward<br>Reverse | CAATTGGAACGCACGAGTCAA<br>GGGCTGTTTGTCCATTACCCA | *tetR21* | 176 |
| OLUG1-F<br>OLUG1-R | Forward<br>Reverse | GGTGGCGACTTTGATCTAGC<br>TTATACAACGGTGGCTGTGC | *gyrB* | 169 |
| OLUG2-F<br>OLUG2-R | Forward<br>Reverse | TTTACGTGCAGCACGTTCAC<br>AAAAAGAAGCTGGTTCAGCAGTAG | *hU* | 125 |
| Pr1-F<br>Pr1-R | Forward<br>Reverse | TATGACGTCCACGCGTGGTTGTGTT TGCGTCACG<br>CCCGGGAGCCGAATTCAAT CATCTACACCAATGACAGTGC | *tet38 upstream* | 981 |
| Pr2-F<br>Pr2-R | Forward<br>Reverse | CTCCCGGGTACCATGGATAGAT ATAAATTCGCGAGAT<br>TTAACTAGACAGATCTCTCAAT TTAGTATAGTATGCTTCTG | *tet38 dowstream* | 1000 |
| OPB40F<br>OPB40R | Forward<br>Reverse | TATCGCCGTTTGGTGGTACG<br>GTCACACCCGGCATTACCAT | *norA* | 178 |
| OPB41F<br>OPB41R | Forward<br>Reverse | TGGTCAATTGGCTCATGGGG<br>ACGCCAACCTAAAAGTGTTGC | *norB* | 78 |
| OPB42bF<br>OPB42bR | Forward<br>Reverse | AGTTGGCGTTGCTTCAGGTA<br>CCGGCATATACTGCACCACT | *norC* | 87 |
| OPB21bF<br>OPB21bR | Forward<br>Reverse | CTGGAACGATGGAATGGGCT<br>ACGTACCGCAAATACTGCAA | *sstA* | 114 |
| OPB48F<br>OPB48R | Forward<br>Reverse | GGGCATTAGATGCGACAGCA<br>ACTTCGATCTTTGCGCTTGG | *sak* | 70 |
| OP45F<br>OP45R | Forward<br>Reverse | ATGGCCAGAGTTACGAGTGT<br>CGTCATTTTCCCTTGACCAGC | *nrdF* | 93 |
| OPB49F<br>OPB49R | Forward<br>Reverse | CCACTGCACTCGCACGATTA<br>TTGCTAGTTTTATCATTGGGAGCA | *scn_3* | 70 |
| HlaF<br>HlaR | Forward<br>Reverse | GGTAATGTTACTGGTGATGATACAGGAA<br>TGCAAATGTTTCGATTGGTCATACAC | *hla* | 72 |
| saeR-F1<br>saeR-R1 | Forward<br>Reverse | TGACCCACTTACTGATCGTGG<br>ACCGCTAGTTGTCGTTGTTACT | *saeR* | 154 |
| OBP46F<br>OPB46R | Forward<br>Reverse | GGGCTAAGTCTATTAGGTGGCG<br>ACGTGCTCGAAATGCTTTGAC | *nrdD* | 78 |
| OPB52F<br>OPB52R | Forward<br>Reverse | TCACCAGCAGCATTAGCGAT<br>TGCTTTGACCGCATCACTCT | *aur* | 93 |

**Table S3: MIC of the 12 couples of strains**

| | MIC (µg/ml) | |
|---|---|---|
| Strains | Tetracycline | Ciprofloxacin |
| SA27 | 0,7 | 8 |
| SA30 | 11,25 | 1 |
| SA31 | 0,5 | 8 |
| SA42 | 1 | 1 |
| SA69 | 0,5 | 4 |
| SA80 | 8 | 2 |
| SA82 | 0,5 | 2 |
| SA146 | 0,5 | 4 |
| SA152 | 1 | 16 |
| SA153 | 1 | 8 |
| SA156 | 4 | 2 |
| SA2599 | 4 | 4 |
| PA27 | 8 | 2 |
| PA30 | 8 | 1,5 |
| PA31 | 12 | 0,5 |
| PA42 | 4 | 0,047 |
| PA69 | 1,5 | 1,5 |
| PA80 | 4 | 0,38 |
| PA82 | 8 | 8 |
| PA146 | 24 | 0,38 |
| PA152 | 16 | 1 |
| PA153 | 12 | 2 |
| PA156 | 8 | 0,19 |
| PA2600 | 16 | 6 |

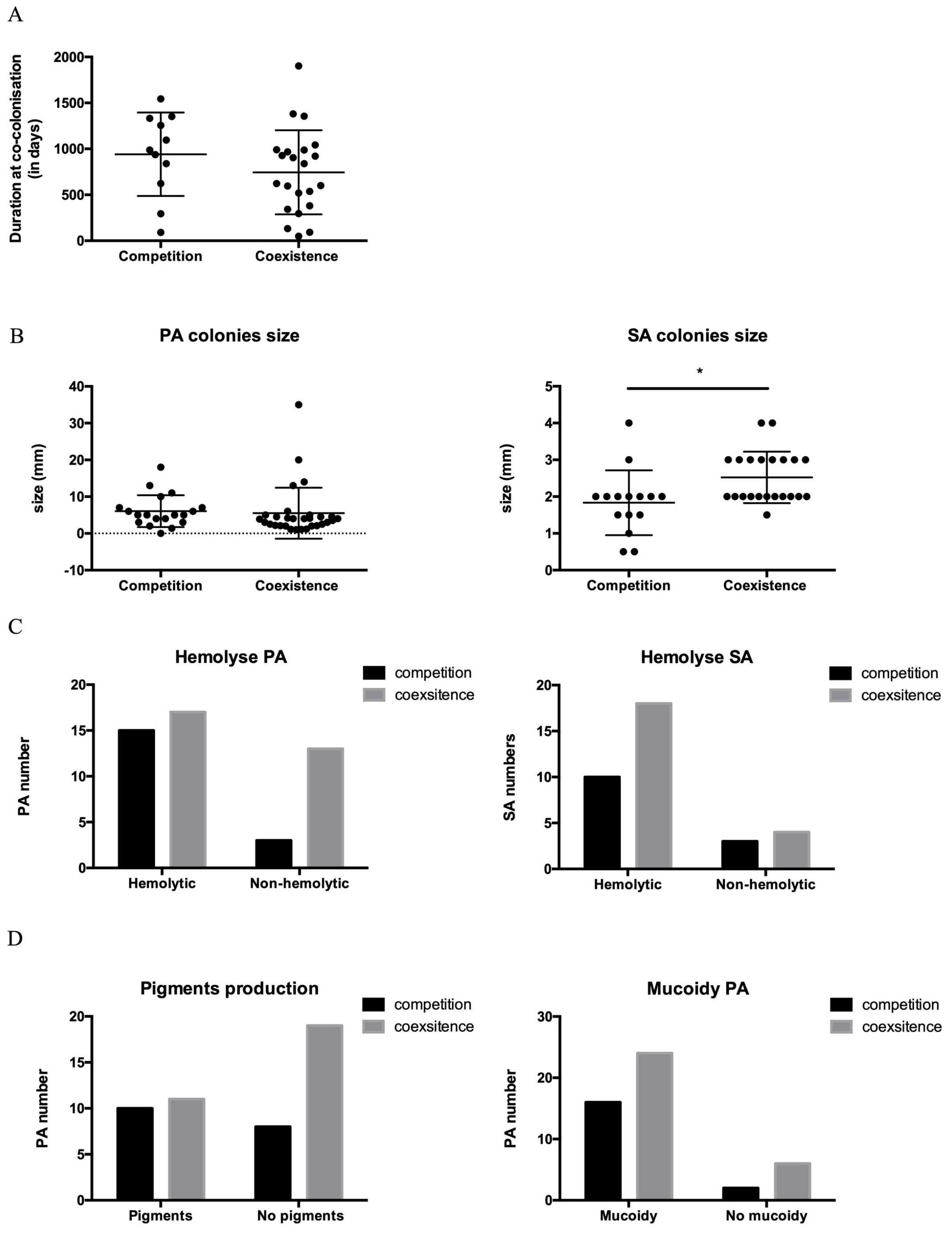


**Figure S1**: Statistical analysis of phenotypic differences between coexisting and competitive strains. No significant differences were observed for the duration of co-colonization (**A** – T-test), *P. aeruginos*a colonies size (**B**- T test), hemolytic properties (**C** – Fisher test), *P. aeruginosa* pigment production and mucoid phenotype (**D**- Fisher test). A significant difference was observed for *S. aureus* colony sizes (**B**- T test, * P<0.05).

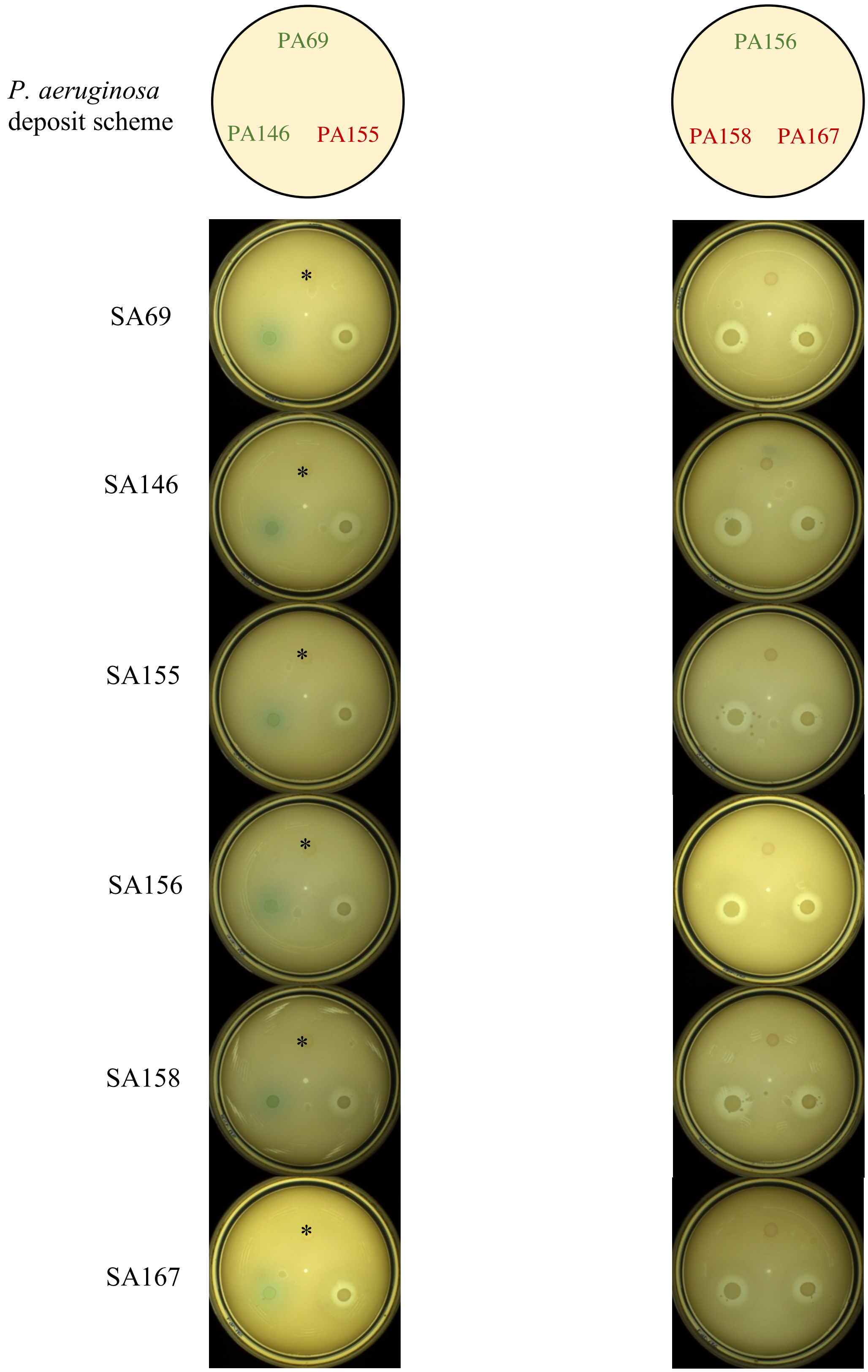


**Figure S2:** Crossed agar competition assay. *S. aureus* - *P. aeruginosa* strain pairs 69, 146, 155, 156, 158 and 167 were cultivated over-night in BHI at 37°C. *S. aureus* strains were plated at $OD_{600nm}$=0.5 onto TSA as indicated on the left of the photograph and left to dry for 15min. *P. aeruginosa* spots were deposited as depicted in the top scheme. *P. aeruginosa* green tags represent a coexisting isolate (absence of inhibition halo) whereas red tags represent a competition isolate (presence of inhibition halo). * symbols indicate the PA69 spots, which are faintly visible on the pictures.

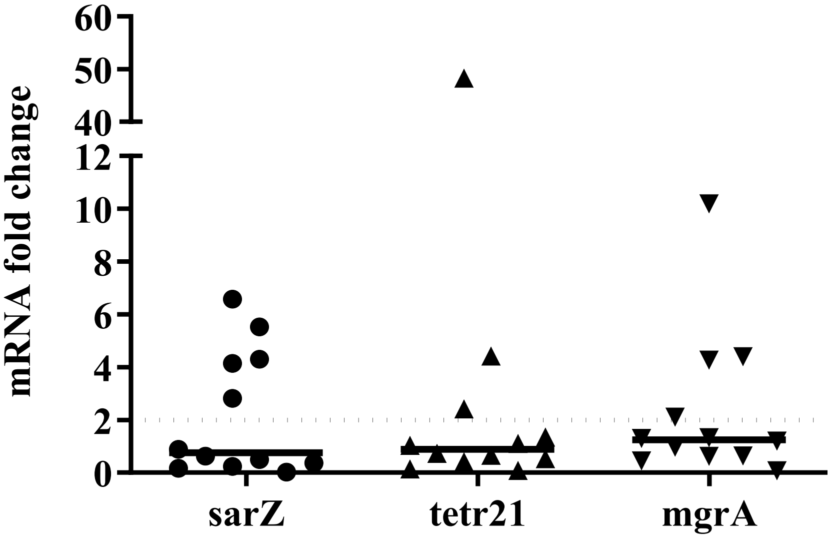


**Figure S3:** Coculture with *P. aeruginosa* has no effect on *tet38* major regulators. RNAs were extracted from a mono- and coculture at 4 hours and gene expression was monitored by RT-qPCR**.** The results represent the median gene expression obtained from twelve coexisting strain pairs. The dotted line depicts fold change= 2.

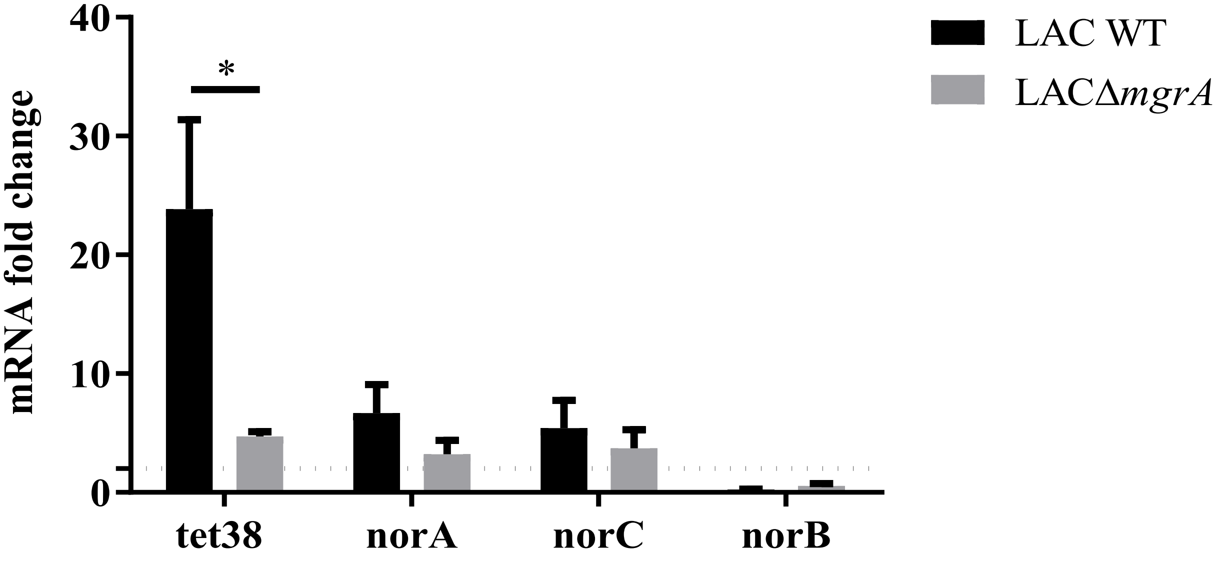


**Figure S4**: MgrA is important for *nor* genes over-expression. Cocultures with *S. aureus* LAC wild type (WT) and *mgrA* mutants (Δ*mgrA*) and PA30 were performed. RNAs were extracted at 8 hours and *nor* gene expression monitored by RT-qPCR. The dotted line indicates a fold change= 2. The results are shown as the mean + standard deviation of three independent experiments. Statistical analysis was performed by unpaired t-test (* P< 0.05)

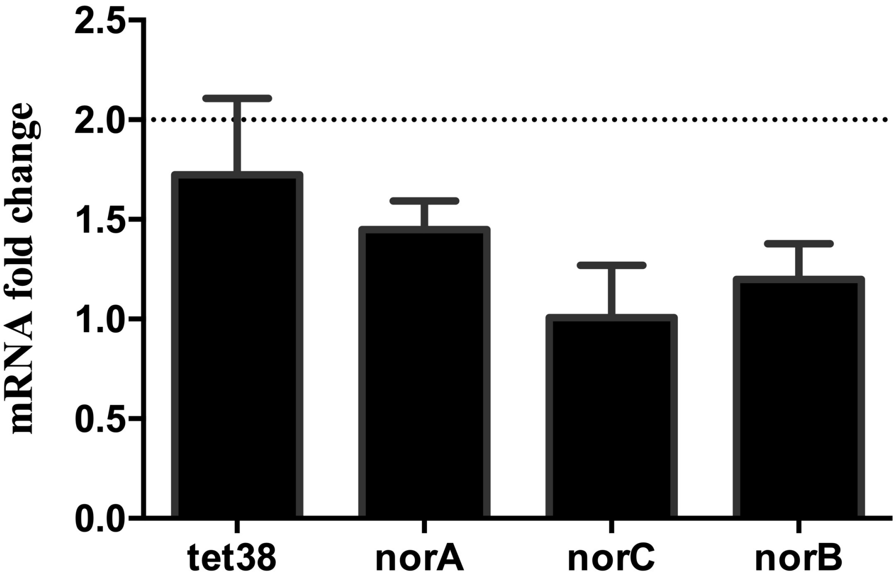


**Figure S5**: *S. aureus nor* genes over-expression requires close contact with *P. aeruginosa*. *S. aureus* was exposed to an 8 hour supernatant culture of *P. aeruginosa*. RNAs were extracted and *nor* gene expression was monitored by RT-qPCR. Dotted lines represent fold change= 2. The results are shown as the mean + standard deviation of three independent experiments on SA30-PA30 pairs. Statistical analysis was performed by unpaired t-test (* $P<0.05$, ** $P<0.01$).

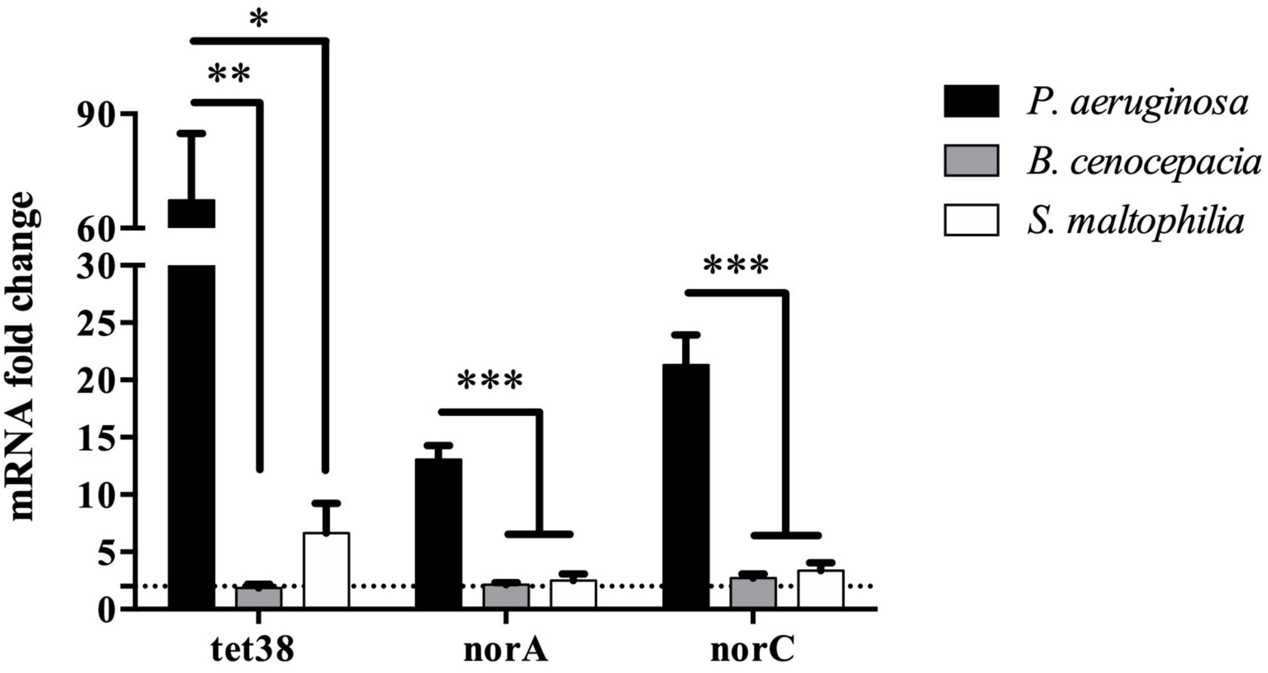


**Figure S6**: Specificity of *nor* genes family over-expression induced by *P. aeruginosa*. *S. aureus* was mono- and co-cultivated as described earlier. RNAs were extracted at 8 hours and gene expression was monitored by RT-qPCR. The dotted line indicates a fold change= 2. The results represent the mean + standard deviation of three independent experiments conducted with pair SA146-PA146. Statistical analysis was performed by One-way Anova with the Dunnett correction multiple test (* P< 0,05, ** P < 0,01, *** P<0,001).

A

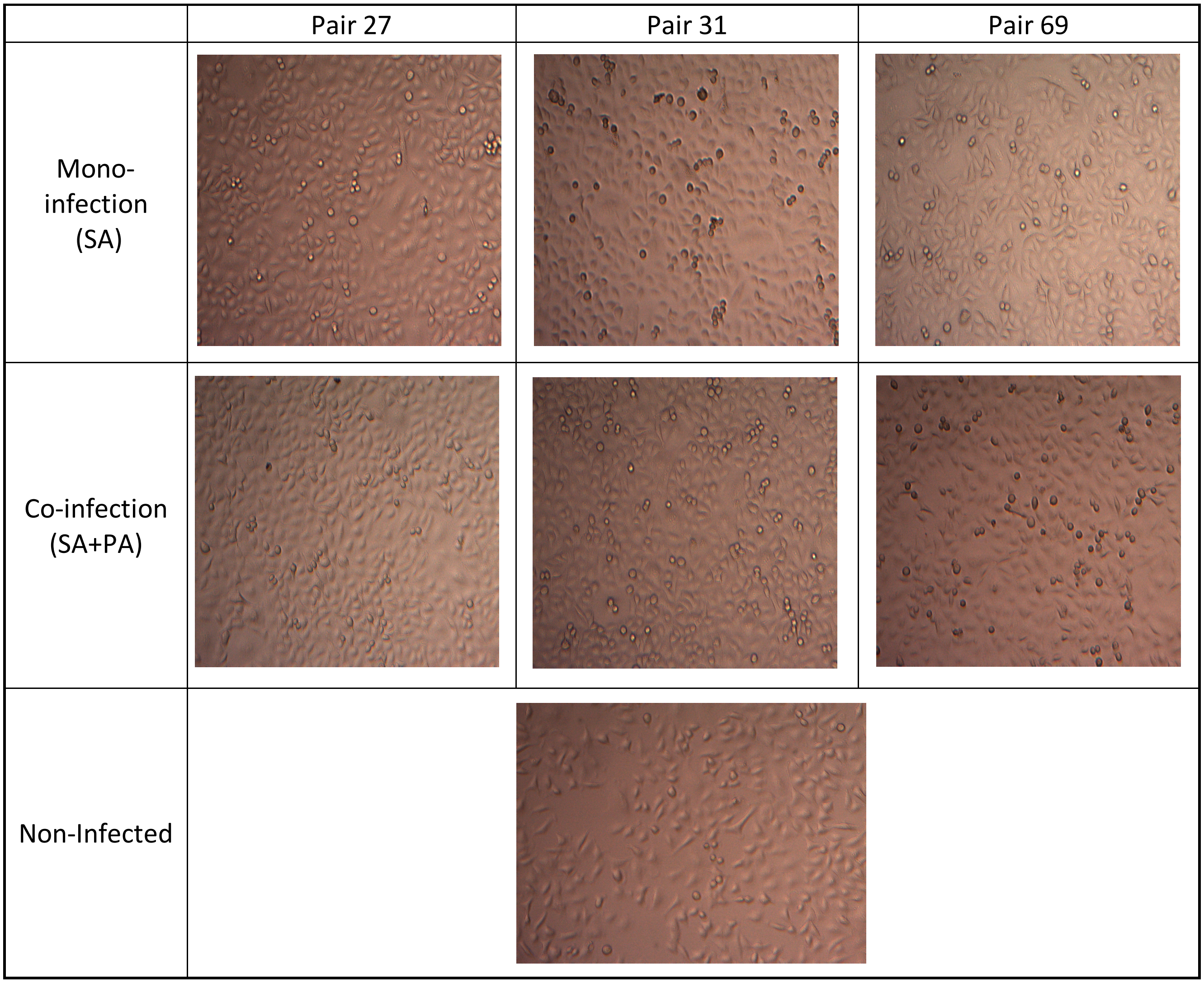

Pair 27
Pair 31
Pair 69
Mono-infection (SA)
Co-infection (SA+PA)
Non-Infected


B

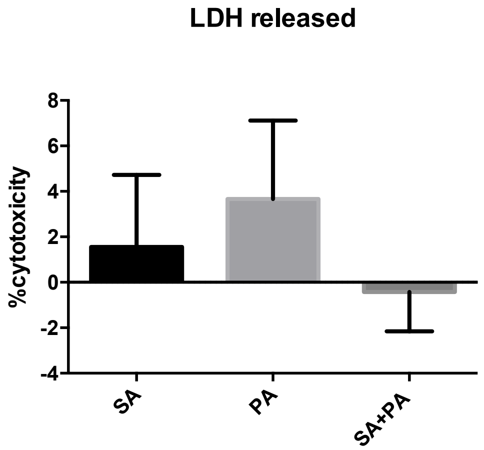

LDH released
%cytotoxicity
8
6
4
2
0
-2
-4
SA
PA
SA+PA

**Figure S7**: Co-infection with *S. aureus* and *P. aeruginosa* do not impact cell shape and viability. A549 cells were infected at a multiplicity of infection (MOI) of 10:1 for mono-culture and 20:1 for coculture. After 2 hours of infection and 1 hour of antibiotic treatment to eliminate extracellular bacteria, cells were photographed (**A**) and LDH was measured in the supernatant using CytoTox-ONE™ Homogeneous Membrane Integrity Assay (Promega) (**B**) following the manufacturer's instructions.
